# Hyperglycosylation of prosaposin in tumor DCs promotes immune escape in cancer

**DOI:** 10.1101/2023.06.14.545005

**Authors:** Pankaj Sharma, Xiaolong Zhang, Kevin Ly, Ji Hyung Kim, Qi Wan, Jessica Kim, Mumeng Lou, Lisa Kain, Luc Teyton, Florian Winau

## Abstract

Tumors develop strategies to evade immunity by suppressing antigen presentation. Here, we show that prosaposin drives CD8 T cell-mediated tumor immunity and that its hyperglycosylation in tumor DCs leads to cancer immune escape. We found that lysosomal prosaposin and its single saposin cognates mediated disintegration of tumor cell-derived apoptotic bodies to facilitate presentation of membrane-associated antigen and T cell activation. In the tumor microenvironment, TGF-β induced hyperglycosylation of prosaposin and its subsequent secretion, which ultimately caused depletion of lysosomal saposins. In melanoma patients, we found similar prosaposin hyperglycosylation in tumor-associated DCs, and reconstitution with prosaposin rescued activation of tumor-infiltrating T cells. Targeting tumor DCs with recombinant prosaposin triggered cancer protection and enhanced immune checkpoint therapy. Our studies demonstrate a critical function of prosaposin in tumor immunity and escape and introduce a novel principle of prosaposin-based cancer immunotherapy.

**One Sentence Summary:** Prosaposin facilitates antigen cross-presentation and tumor immunity and its hyperglycosylation leads to immune evasion.

---

Antigen-specific T cell responses are central to protection against cancer. Cytotoxic CD8 T lymphocytes recognize tumor antigens presented by MHC-I molecules and subsequently deploy their effector functions, such as target cell killing and production of inflammatory cytokines (*1*). However, tumor cells often fail to directly activate T cells due to downregulation of their MHC-I pathway (*2, 3*). Therefore, other antigen-presenting cells, such as dendritic cells (DCs), are critical to engulf tumor antigens for subsequent processing and display to MHC-I-restricted CD8 T cells in a process called cross-presentation (*4*). On a cellular level, this mechanism can be broadly divided into a cytosolic and vacuolar pathway (*5, 6*). According to cytosolic processing, endosomal antigens are retrotranslocated into the cytosol for degradation by proteasomes and subsequent reimport into the endosome for MHC-I loading. By contrast, in the vacuolar pathway, the endosome is more autonomous and relies on its proteases for antigen processing (*7–9*). Among the different DC subsets, classical DC1 are especially efficient in cross-presentation and fulfill this function in tumor-draining lymph nodes for T cell priming as well as in the tumor microenvironment to activate tumor-infiltrating T lymphocytes (*10, 11*). Abundance and proper function of immune cells, including antigen-presenting cells and lymphocytes, at the tumor site are vital for effective immunity and control of cancer growth (*12, 13*). Unfortunately, tumors often develop mechanisms to evade immune responses, for example by producing immunosuppressive cytokines, such as TGF-β (*14, 15*). Thus, the goal of cancer therapy is to target and overcome immune evasion to restore protective immunity.

Prosaposin is a precursor protein that is transported from the Golgi apparatus to the lysosome assisted by its chaperone sortilin (*16, 17*). In the lysosome, cathepsins cleave prosaposin into the single saposins A-D. Saposins are also called sphingolipid activator proteins since they function as small, non-enzymatic cofactors for lysosomal hydrolases that are required for sphingolipid degradation (*18*). Moreover, at the low acidic pH of the endolysosomal compartment, saposins are able to interact with anionic phospholipids, such as phosphatidylserine (PS), exposed on intralysosomal vesicles (*19*). These membrane-perturbing properties facilitate vesicle disintegration and also pertain to apoptotic vesicles that characteristically contain PS in their lipid bilayers (*20, 21*). In this context, tumors, owing to their uncontrolled growth kinetics, produce a substantial amount of dying cells and apoptotic bodies, which contain tumor antigens to potentially trigger the immune system. Notably, membrane-associated particulate antigen is more immunogenic than soluble protein and thus, antigen presentation pathways based on vesicular processing might be central to the induction of protective T cell immunity. In this study, we explored the impact of saposins on presentation of membrane-associated tumor antigen and activation of CD8 T cell responses that protect against cancer growth. This work also describes a mechanism how the tumor counteracts saposin-mediated processing by triggering prosaposin hyperglycosylation and secretion from tumor-associated DCs. Ultimately, we test the proof-of-principle of prosaposin targeting to DCs as a mode of cancer immunotherapy.

## Results

### Saposins promote cross-presentation of membrane-associated tumor antigen

First, we investigated the effect of saposins on the integrity of apoptotic bodies derived from tumor cells. For this purpose, we exposed murine MCA fibrosarcoma cells to γ-irradiation (100 Gy) to trigger apoptotic cell death (Fig. 1A). Successful induction of apoptosis was controlled by measuring phosphatidylserine exposure using AnnexinV staining. Subsequently, we purified apoptotic vesicles from cell culture supernatants using differential ultracentrifugation (100,000 *g* pellets) and visualized them by transmission electron microscopy (Fig. 1B). We then loaded the fluorescent dye calcein into those apoptotic vesicles using a liposome extruder with a 100 nm pore size. After incubation with different recombinant saposins, we measured calcein release and found that saposins disintegrate tumor cell-derived apoptotic vesicles when compared to control BSA (Fig. 1B).

**Fig. 1.**
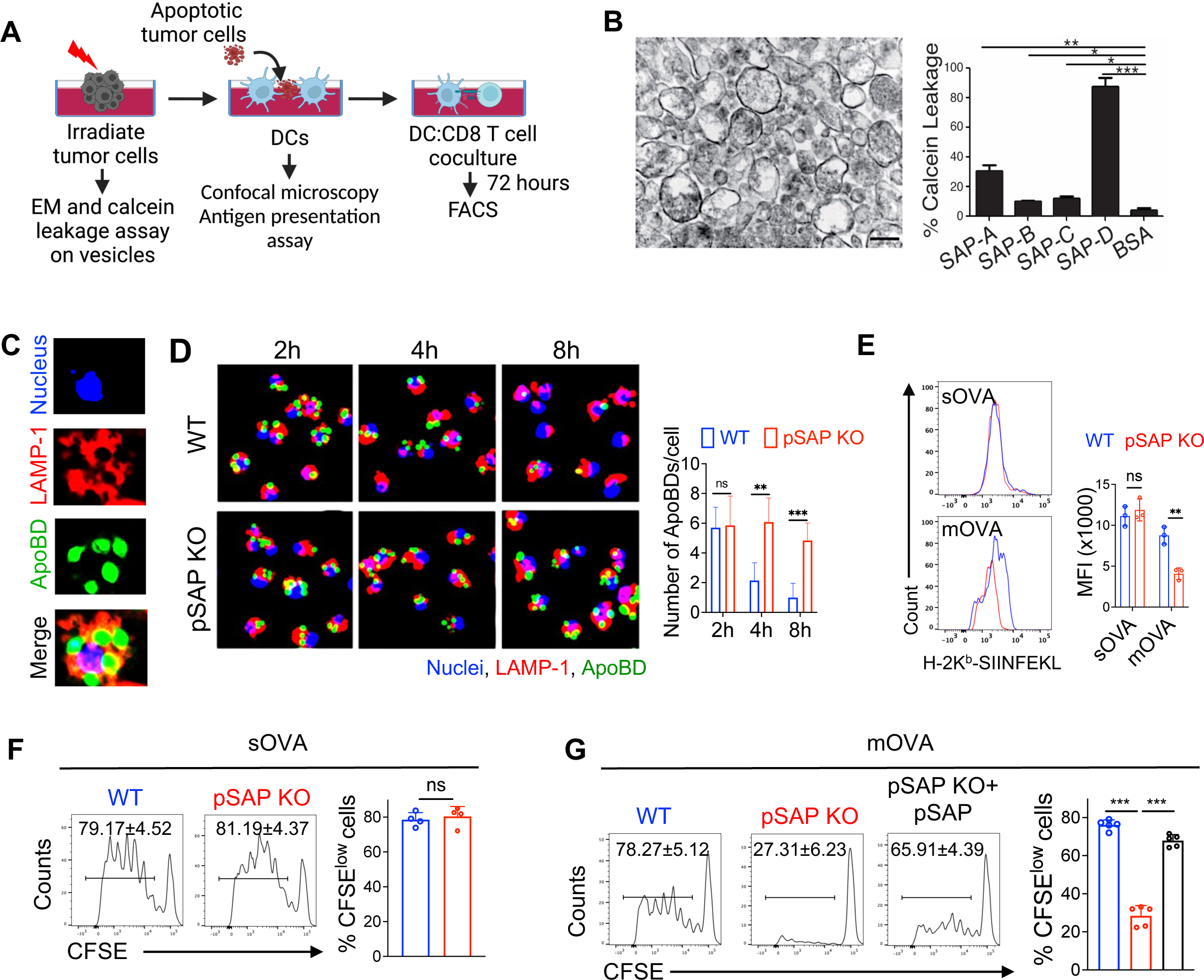
Saposins promote cross-presentation of membrane-associated tumor antigen. **A.** Diagram depicting the experimental read-outs used in Figure 1. MCA101 fibrosarcoma cells were γ-irradiated (100 Gy), prior to collection of apoptotic vesicles from supernatant and analysis using electron microscopy (EM) and calcein leakage assay. Moreover, apoptotic MCA101 cells expressing membrane-associated ovalbumin were used to pulse bone marrow-derived DCs from WT or pSAP-KO mice, prior to analysis of digestion of apoptotic cells using confocal microscopy, and antigen processing and T cell activation using FACS. **B.** Calcein leakage assay to quantify the effect of saposins on disintegration of apoptotic bodies. Apoptotic vesicles were prepared using differential ultracentrifugation (100,000 *g*) from supernatant of irradiated MCA101 cells and visualized using transmission electron microscopy (left image). Scale bar = 200 nm. Apoptotic bodies were further loaded with calcein dye prior to incubation with indicated saposins or BSA (negative control), and calcein release was quantified in the supernatant using fluorimetry. Depicted is % leakage compared to 100% lysis induced by Triton X-100. **C.** Representative confocal microscopy image showing colocalization of apoptotic bodies (green) with LAMP-1 (red). WT DCs were pulsed with CFSE-labeled, γ-irradiated apoptotic MCA101 tumor cells for 2 hours. ApoBD = apoptotic body. **D.** Representative confocal microscopy images showing the kinetics of apoptotic cell disintegration in WT or pSAP-KO DCs. DCs were pulsed with CFSE-labeled, γ- irradiated apoptotic MCA101 tumor cells for 2 hours, and the numbers of apoptotic bodies (ApoBD) were quantified at indicated time points using ImageJ software (DAPI: blue, LAMP-1: red, apoptotic bodies: green). **E**. Representative histogram overlays and bar graph showing flow cytometry staining and mean fluorescence intensity (MFI) of MHC-I-SIINFEKL peptide on the surface of WT or pSAP-KO DCs after incubation with either soluble OVA (sOVA) or irradiated MCA101-OVA tumor cells (mOVA = membrane-associated OVA) for 4 hours. **F**. Histograms and bar graph showing frequencies of proliferating, CFSE^low^ CD8 T cells after 3-day coculture with WT or pSAP-KO DCs pulsed with soluble OVA. **G.** Histograms and bar graph depicting the frequencies of CFSE^low^ CD8 T cells after 3-day coculture with WT or pSAP-KO DCs pulsed with irradiated MCA101-OVA cells. In addition, pSAP-KO DCs were reconstituted with 10 µg/ml of recombinant prosaposin prior to the T cell assay. Data shown in all graphs represent mean ±SD from 3-5 independent replicates. *p*-values were determined using unpaired Student’s t-test. *p < 0.05; **p < 0.01; ***p < 0.001; ns: not significant.

We next explored the impact of saposins on processing of apoptotic bodies in DCs. To this end, we pulsed bone marrow-derived DCs with CFSE-labeled apoptotic cells derived from γ-irradiated MCA101 tumor cells and followed their fate along the endolysosomal compartment. Confocal microscopy revealed colocalization of apoptotic bodies with LAMP-1, indicating trafficking to saposin-containing lysosomes (Fig. 1C). When comparing the kinetic digestion of apoptotic cells in prosaposin-deficient or wild-type DCs, we found that early uptake of apoptotic material was similar, suggesting that phagocytosis is not affected by saposin deficiency. However, at later time points after endocytosis, prosaposin-deficient DCs accumulated CFSE-labeled cells, demonstrating the importance of saposins for processing of apoptotic bodies in DCs (Fig. 1D).

To test for saposin-dependent cross-presentation and CD8 T cell activation, we pulsed DCs from prosaposin-KO or WT mice with apoptotic MCA101 cells expressing a membrane-associated form of the antigen ovalbumin (OVA), prior to coculture with OVA-specific CD8 T cells. First, we analyzed productive antigen processing by staining for the processed OVA epitope in complex with MHC-I (H-2k^b^-SIINFEKL) on the DC surface. Flow cytometry demonstrated that processing of soluble OVA was equally efficient in pSAP-KO and WT DCs (Fig. 1E). However, processing and loading onto MHC-I of membrane-associated antigen derived from tumor cells was significantly hampered in the absence of prosaposin (Fig. 1E). These findings were in accordance with our T cell data, since pSAP-deficient DCs activated CD8 T cells responding to soluble OVA as efficient as WT DCs (Fig. 1F). In sharp contrast, when DCs were pulsed with tumor cells containing membrane-associated antigen, CD8 T cell activation by pSAP-KO DCs was strikingly reduced, as highlighted by their truncated CFSE dilution profile in flow cytometry (Fig. 1G). Notably, incubation of pSAP-deficient DCs with recombinant prosaposin fully reconstituted CD8 T cell activation (Fig. 1G). Taken together, saposins disintegrate apoptotic vesicles and process membrane-associated antigen for cross-presentation to CD8 T cells.

### Prosaposin is required for tumor immunity

Next, we investigated how prosaposin function affects T cell activation in vivo and protection against cancer. In order to assess T cell priming, we transferred naïve CFSE-labeled, OVA-specific CD8 T cells into pSAP-deficient or WT recipients, prior to subcutaneous administration of apoptotic MCA101-OVA cells (fig. S1A). Four days after tumor cell injection, we isolated DCs from skin-draining lymph nodes and analyzed antigen processing. Expression of H-2k^b^-SIINFEKL proved to be reduced in DCs from pSAP-KO when compared to WT mice (fig. S1B). Moreover, antigen-specific proliferation and IFN-γ production by CD8 T cells from lymph nodes were severely hampered in the absence of prosaposin (fig. S1C). Thus, prosaposin facilitates CD8 T cell priming in response to particulate antigen. Since straight pSAP-KO mice have a reduced life span, we then generated chimeric mice by transferring pSAP-KO or WT bone marrow to WT recipients in order to allow for tumor challenge experiments. In this context, we immunized mice with irradiated tumor cells and challenged them with a higher number of live MCA101-OVA cells one week later (Fig. 2A). Subsequently, we monitored tumor growth in the skin and found a drastic expansion of cancer in prosaposin deficiency (Fig. 2B). Flow cytometry analysis of isolated DCs from the tumor site showed pSAP-dependent decrease in antigen processing and presentation (Fig. 2C). Furthermore, MHC-I tetramer-mediated detection of antigen-specific CD8 T cells showed reduced frequency of tumor-infiltrating T cells as well as cytokine production in prosaposin-deficient mice (Fig. 2D). In addition, we also challenged bone marrow-chimeric mice with live tumor cells without prior vaccination (fig. S2A). As a result, cancer protection, antigen processing in tumor DCs, frequency of tumor-infiltrating, antigen-specific T cells, as well as cytokine production and cytotoxicity were all strikingly reduced when prosaposin was lacking (fig. S2B-E). Altogether, these findings demonstrate that tumor immunity critically depends on prosaposin function.

**Fig. 2.**
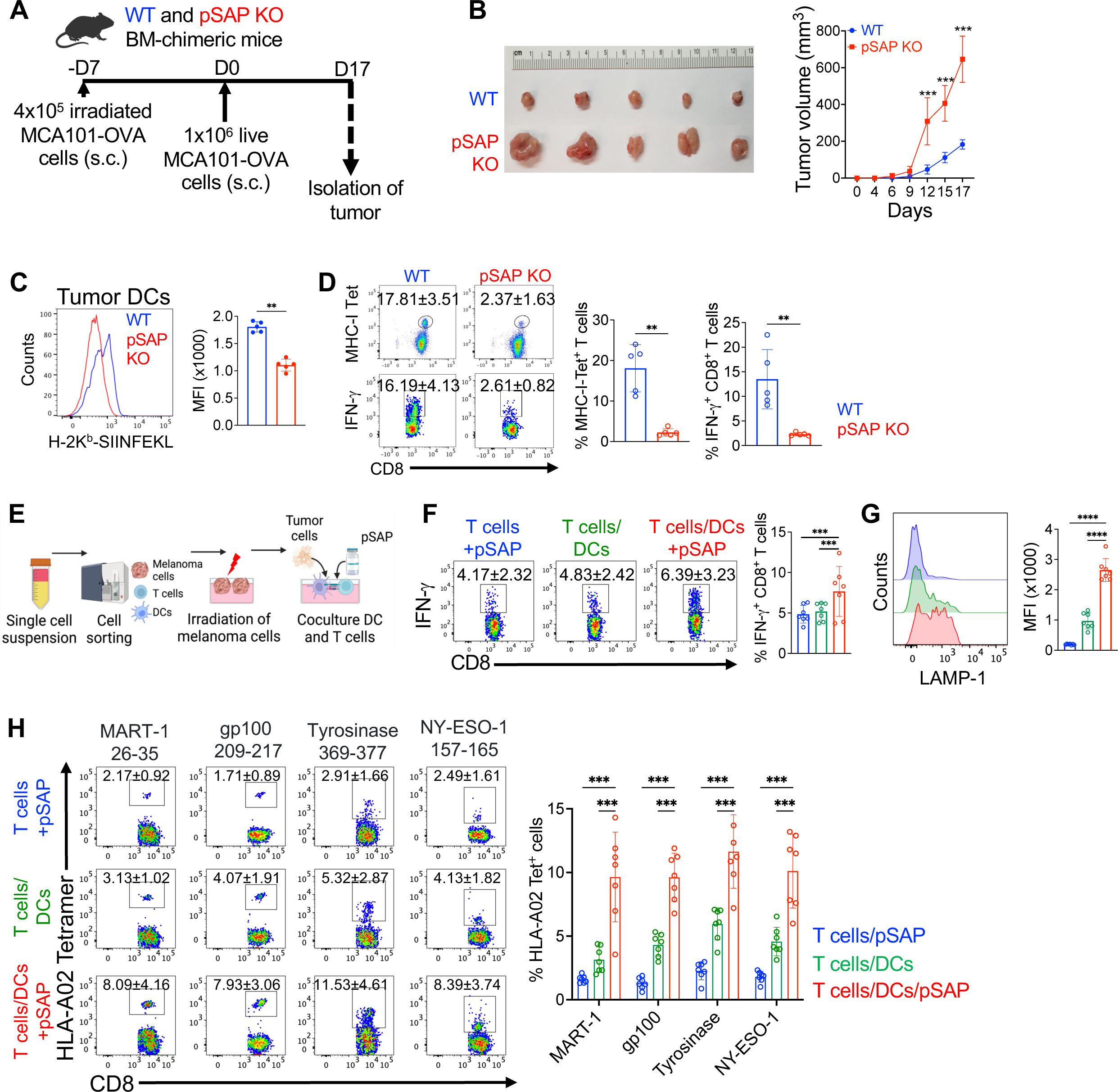
Prosaposin is required for tumor immunity and boosts T cells from melanoma patients. **A.** Experimental scheme of tumor challenge studies. WT and pSAP-KO BM chimeric mice were primed with 4 x 10^5^ γ-irradiated MCA101-OVA cells (s.c.) and subsequently inoculated with 1 x 10^6^ live MCA101-OVA cells (s.c.) 7 days post priming. **B**. Comparison of tumor sizes between WT and pSAP-KO mice on day 17 (left) and kinetics of tumor growth (right). **C**. Representative histogram overlay and bar graph depicting the staining and mean fluorescence intensity (MFI) of MHC-I-SIINFEKL peptide on the surface of tumor DCs from pSAP-KO or WT animals. **D**. FACS plots and bar graphs showing frequencies of MHC-I (Kb-SIINFEKL) tetramer- and IFN-γ-positive tumor-infiltrating CD8 T cells in pSAP-KO or WT mice. MHC-I tetramer specifically detects CD8 T cells reactive with SIINFEKL peptide. **E**. Experimental set-up for the coculture of myeloid and CD8 T cells isolated from human melanoma. Single cell suspensions from human melanoma samples were FACS-sorted for CD146^+^ melanoma cells, CD8^+^ T cells, and CD11c/b^+^ myeloid cells. CD146^+^ cells were γ-irradiated and incubated with DCs, which were further cocultured with CD8 T cells in the presence or absence of recombinant pSAP. **F.** FACS plots and bar graph showing the frequencies of IFN-γ-positive CD8 T cells following the indicated culture conditions. **G**. Representative histogram overlay and bar graph demonstrating surface staining and MFI of LAMP-1 on CD8 T cells according to the indicated culture conditions. **H**. Flow cytometry analysis and summarizing bar graph depicting the frequencies of antigen-specific CD8 T cells reactive with HLA-A*0201 tetramers loaded with epitopes from gp100, MART-1, Tyrosinase, and NY-ESO-1, following the indicated culture set-ups. Data shown in graphs B-D are representative of three independent experiments, while F-H depict mean ±SD from 7 independent subjects. *p*-values were determined by unpaired Student’s t-test in graphs B-D, while one-way ANOVA was used in graphs F-H. **p < 0.01; ***p < 0.001; ****p < 0.0001.

### T lymphocytes from melanoma patients are boosted by prosaposin

To study prosaposin in the context of human cancer, we used dissociated tumor cell (DTC) samples from melanoma patients. A detailed description of the patient samples can be found in table S1. Briefly, the majority of specimens were isolated from primary melanoma lesions from the skin, were assigned a clinical stage III, before treatment, including white, female and male patients older than 50 years (table S1). To purify antigen-presenting cells, responder T cells, and tumor cells as source of antigen, we FACS-sorted CD146^+^ melanoma cells, CD11b/c^+^ myeloid cells, and CD8^+^ T cells (fig. S3). We then irradiated CD146^+^ melanoma cells and pulsed them onto sorted myeloid cells, prior to coculture with autologous CD8 T cells (Fig. 2E). In parallel, we also treated DCs and T cells with human recombinant prosaposin. Five days after culture, we analyzed effector functions of CD8 T cells and found that recombinant prosaposin was able to boost IFN-γ production (Fig. 2F), as well as cytolytic activity as indicated by surface LAMP-1 staining as sign of cytotoxic degranulation (Fig. 2G). Furthermore, we measured the frequencies of tumor antigen-specific CD8 T cells by staining with MHC-I tetramers loaded with dominant melanoma antigens, including MART, gp100, Tyrosinase, and NY-ESO-1. The abundance of melanoma-specific CD8 T cells was strikingly increased when tumor DCs were treated with prosaposin (Fig. 2H). Of note, the patient samples have been HLA-typed by flow cytometry beforehand to select the proper haplotype for MHC-I tetramer analysis (HLA-A02). To conclude, the impact of prosaposin on tumor DCs is able to rescue T cell activation from melanoma patients.

### Hyperglycosylation of prosaposin in tumor DCs leads to its secretion

Beyond the use of pSAP deficiency in a mouse model, we aimed at understanding the regulation of prosaposin in tumor-associated DCs in a pathophysiological context. To this end, we inoculated WT mice with live MCA101-OVA cells subcutaneously and subsequent to tumor outgrowth, we isolated DCs from the tumor microenvironment (TME), tumor-draining lymph nodes, and spleen (fig. S4A). We FACS-purified the two main classical DC subsets based on their established markers as cDC1 (XCR1^+^ or CD103^+^) and cDC2 (SIRP1α^+^ or CD11b^+^), prior to performing an array of antigen processing and presentation assays. Accordingly, pulsing with FITC-dextran showed that the phagocytosis rate of tumor DCs was not altered when compared to lymph node and spleen (fig. S4B). Incubation with a self-quenched antigen conjugate (DQ-OVA), which exhibits fluorescence upon proteolytic degradation, demonstrated that mainly cDC2 in the tumor are compromised to process soluble antigen (fig. S4C). These findings were in line with the ultimate epitope expression on surface MHC-I following pulsing with soluble OVA, which revealed hampered antigen presentation by tumor cDC2 (fig. S4D). In sharp contrast, the presentation capacity of cDC1 in the TME was only affected after incubation with irradiated MCA101-OVA cells, which contain antigen in membrane-associated form (fig. S4D). This phenomenon was reflected in functional T cell experiments, because DCs isolated from tumors were severely perturbed to induce T cell responses reactive to membrane-associated antigen (fig. S4E). Since we found that saposins are critical for presentation of particulate antigen, we then hypothesized that prosaposin function might be modulated in tumor DCs as a basis for poor T cell induction in the TME.

Indeed, analysis by immunoblot revealed the expression of a 75 kDa high-molecular weight form of prosaposin in tumor DCs when compared to pSAP-65 predominant in DCs from spleen (Fig. 3A). Moreover, the small, single saposins were severely depleted in DCs from the TME (Fig. 3A). When we cultured the respective DC subsets ex vivo, we observed secretion of prosaposin into the cell culture supernatant mainly by tumor DCs as measured by ELISA (Fig. 3B). In order to demonstrate that the occurrence of pSAP-75 was due to glycosylation, we tested endoglycosidase H (Endo H) sensitivity of prosaposin. Endo H cleaves N-linked glycans between the two proximal N-acetylglucosamine residues only in high-mannose carbohydrate chains, but not in complex glycans. After treatment of protein lysates with Endo H, pSAP-65 from splenic DCs was cleaved to lower molecular weight forms, whereas pSAP-75 from tumor DCs proved to be endo H-resistant, suggesting that it contained complex glycans (Fig. 3C). To corroborate this finding, we performed deeper molecular analysis using mass spectrometry of sugar structures based on purified pSAP bands derived from splenic (pSAP-65) or tumor DCs (pSAP-75). Tandem mass spectrometry showed that the glycan of pSAP-65 mainly consisted of mannose residues, while pSAP-75 exhibited complex glycans involving additions of N-acetylglucosamine, galactose, and sialic acid (Fig. 3D). Since glycan structures are synthesized by a diverse set of glycosyltransferases, we compared the expression of glycosyltransferases between tumor and splenic DCs using a qRT-PCR array (Fig. 3E). In tumor DCs, we found upregulation of several enzymes that facilitate the attachment of complex glycan residues, such as N acetylglucosaminyltransferases, galactosyltransferases, and sialyltransferases (Fig. 3F).

**Fig. 3.**
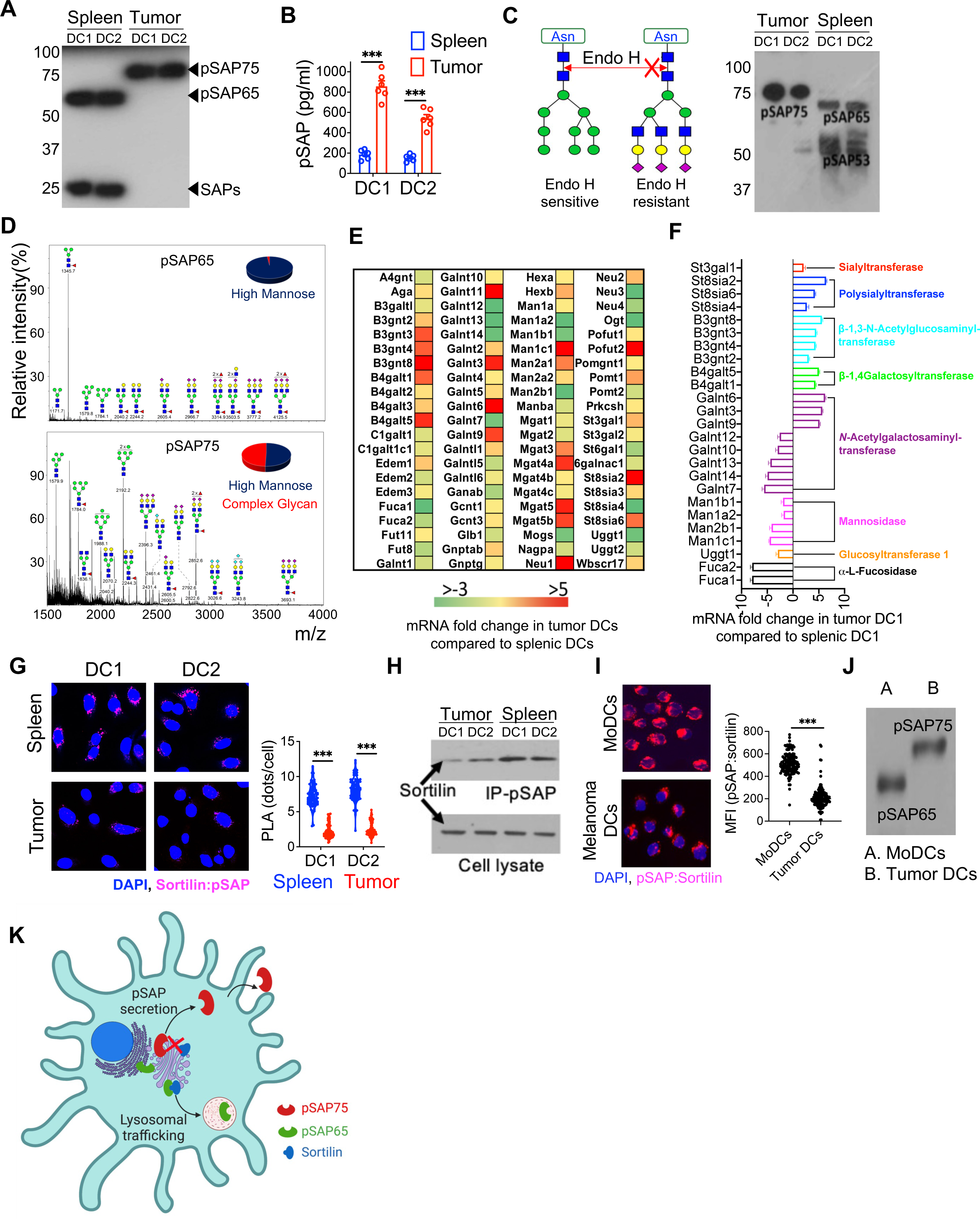
Hyperglycosylation of prosaposin in tumor DCs leads to its secretion. WT mice were inoculated with 1 x 10^6^ live MCA101 cells, and 18 days post tumor inoculation, cDC1 and cDC2 populations were FACS-sorted from tumor and spleen. **A.** Immunoblot showing the expression of pSAP and saposins in tumor and splenic DC subsets. pSAP-75 = hyperglycosylated prosaposin, pSAP-65 = glycosylated pSAP, and SAPs = saposins. **B.** Quantification of prosaposin secreted by DCs. FACS-sorted splenic and tumor DC subsets were cultured in cRPMI for 48 hours, and pSAP in culture supernatant was quantified using ELISA. **C**. Immunoblot of Endo H-treated pSAP. Left: Mechanism of Endo H that leads to cleavage of high-mannose but not complex glycans. Right: FACS-sorted DCs from tumor and spleen were lysed in RIPA buffer, and cell lysates were treated with Endo H for 12 hours at 37°C, prior to analysis using immunoblot. **D**. MALDI-TOF mass spectrometry analysis of permethylated N-linked glycans of prosaposin immunoprecipitated from FACS-sorted CD11c^+^ DCs. Enzymatically released N-glycans from pSAP of splenic (top panel) and tumor (bottom panel) DCs were analyzed. Glycan compositions were assigned based on m/z values. x-axis: mass to charge ratio (m/z). y-axis: signal intensity of the ions. green circle = mannose; yellow circle = galactose; red triangle = fucose; blue square = N-acetylglucosamine; magenta diamond = sialic acid. **E**. Heat map of differentially expressed genes in tumor DCs involved in glycosylation, as analyzed by real-time RT^2^ profiler PCR array. Splenic DCs were used as control to calculate fold change in gene expression. **F**. Bar graph depicting glycosyltransferase and glycosidase gene expression in tumor compared to splenic DC1. **G**. Proximity ligation assay (PLA) of pSAP and sortilin. Confocal microscopy images of tumor and splenic DC subsets reveal PLA signal between pSAP and sortilin. Blue indicates cell nucleus (DAPI), and magenta represents ligation signal. The violin plot shows quantification of PLA signal, where 200 cells from each sample were analyzed for statistics. **H**. Co-immunoprecipitation of sortilin and pSAP in tumor and splenic DCs. Top panel shows blot of sortilin pulled down by anti-pSAP antibody. Bottom panel demonstrates immunoblot of total sortilin in corresponding DC populations. **I**. PLA of pSAP and sortilin in human melanoma and monocyte-derived DCs. Melanoma DCs were sorted as CD11c^+^ cells from viable CD45^+^ cells isolated from human melanoma samples, while monocyte derived DCs (MoDCs) were generated by culturing monocytes with interleukin 4 (IL-4) and granulocyte-macrophage colony-stimulating factor (GM-CSF) for 4 days. Blue indicates cell nucleus (DAPI), and magenta represents ligation signal. The violin plot shows quantification of PLA signal, where 200 cells from each sample were analyzed for statistics. **J.** Immunoblot of pSAP in human melanoma DCs and MoDCs. pSAP-75 = hyperglycosylated prosaposin; pSAP-65 = glycosylated pSAP. **K.** Illustration visualizing glycosylation mechanisms that control prosaposin trafficking in tumor DCs. Hyperglycosylation of prosaposin compromises its interaction with sortilin and reroutes it to the secretory pathway. Data shown in all graphs are representative of three independent experiments, and *p*-values were determined using unpaired Student’s t-test. *p < 0.05; **p < 0.01; ***p < 0.001; ****p < 0.0001.

Since hyperglycosylation of prosaposin leads to its secretion and therefore reduced generation of single saposins, we next investigated the interaction of pSAP with its chaperone sortilin, which is normally required for efficient lysosomal delivery of prosaposin. For this purpose, we subjected tumor and splenic DCs to a proximity ligation assay (PLA), in which antibodies against pSAP and sortilin are coupled to oligonucleotide probes that allow for subsequent ligation and amplification in case the two target proteins are in close vicinity (10-80 nm). Confocal microscopy revealed these PLA events as discrete spots and showed that the spatial relationship between prosaposin and sortilin was perturbed in DCs from the TME (Fig. 3G). To assess the direct physical interaction between pSAP and sortilin via biochemistry, we performed immunoprecipitation of prosaposin and subsequently resolved sortilin using immunoblot. As a result, the abundance of sortilin recovered from the pSAP precipitate was clearly reduced in tumor DCs, indicating that the interaction of prosaposin with its chaperone sortilin is hampered in the TME (Fig. 3H). To translate these findings to the human system, we explored pSAP/sortilin interaction in DCs isolated from the tumor site of melanoma patients compared to monocyte-derived DCs. The resulting PLA signals demonstrated that the spatial interaction of prosaposin with sortilin was reduced in melanoma DCs (Fig. 3I). Furthermore, we examined the abundance of the different molecular weight forms of prosaposin and found that tumor DCs from melanoma patients exclusively expressed hyperglycosylated pSAP-75 (Fig. 3J). Taken together, our results demonstrate that prosaposin is hyperglycosylated in tumor DCs, fails to interact with its chaperone sortilin, and follows instead a secretory route (Fig. 3K). This mechanism of pSAP hyperglycosylation leads to a depletion of intracellular saposins available for antigen processing, which might explain the compromised antigen presentation capacity in the tumor microenvironment.

### TGF-β induces prosaposin hyperglycosylation in tumor DCs and drives immune evasion

Next, we mechanistically addressed the question how hyperglycosylation of prosaposin is regulated. Considering its prominent immunosuppressive function as a cytokine, we incubated a murine DC line (DC2.4) with recombinant TGF-β and observed a dose-dependent induction of pSAP-75 (Fig. 4A). In addition, treatment with TGF-β was able to trigger secretion of pSAP-75 into the cell culture supernatant as detected by immunoblot and ELISA (Fig. 4A, B). Moreover, we measured gene expression of a panel of enzymes involved in the glycosylation pathway in DC2.4 cells treated with TGF-β (fig. S5). We observed that the upregulated genes in TGF-β- treated DCs correlated well with the enzyme signature detected in tumor DCs (Fig. 4C), suggesting that TGF-β is responsible for triggering the respective glycosylation program. In order to test whether TGF-β signaling is indeed required for prosaposin hyperglycosylation in vivo, we used mice that lack TGF-β receptor II specifically in DCs (CD11c-Cre x Tgfbr2^flox/flox^) for challenge with live MCA101-OVA tumor cells (Fig. 4D). As anticipated, lack of TGF-β downstream signaling in DCs caused better tumor protection, increased antigen presentation in tumor DCs, and stronger IFN-γ production by tumor-infiltrating CD8 T cells (Fig. 4E-G). More importantly, when we isolated tumor DCs for analysis by immunoblot, we found that pSAP-75 was virtually absent in DCs lacking TGF-β signaling (Fig. 4H). Additionally, tumor DCs lacking TGF-β receptor exhibited an abundance of single saposins, which was in striking contrast to the saposin depletion and expression of hyperglycosylated prosaposin predominating over pSAP-65 in WT DCs (Fig. 4H). Furthermore, the enzyme signature involved in glycosylation proved to be reduced when TGF-β signaling was deficient in tumor DCs (Fig. 4I). Thus, TGF-β is essential for hyperglycosylation of prosaposin in tumor DCs, a mechanism associated with immune escape.

**Fig. 4.**
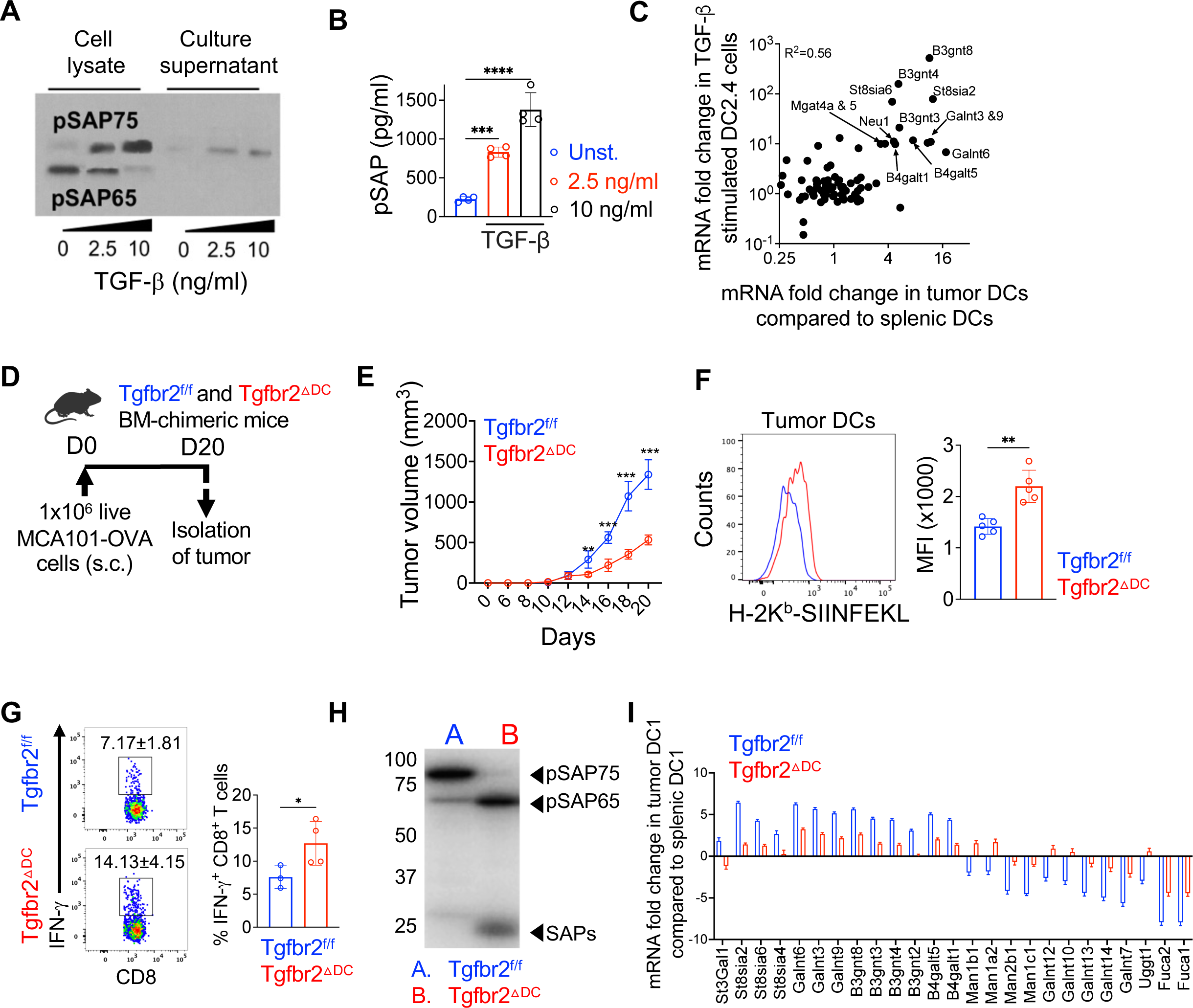
TGF-β induces prosaposin hyperglycosylation in DCs and compromises tumor immunity. **A.** Immunoblot showing dose-dependent induction of pSAP hyperglycosylation and secretion by the murine DC line (DC2.4) incubated with recombinant TGF-β for 48 hours. pSAP-75 = hyperglycosylated prosaposin, pSAP-65 = glycosylated pSAP. **B**. Quantification of pSAP in the culture supernatant of DC2.4 cells after incubation with recombinant TGF-β for 2 days as measured using ELISA. **C**. Scatter plot showing the correlation of gene expression of glycosyltransferases and glycosidases between tumor DCs and TGF-β-stimulated DC2.4 cells. mRNA fold changes were quantified by real-time RT^2^ profiler PCR array. CD11c^+^ tumor DCs were analyzed using splenic DCs as control, while TGF-β-stimulated DC2.4 cells were compared with sham-treated DC2.4 cells. **D**. Experimental scheme of tumor cell challenge. Tgfbr2^f/f^ and CD11c Cre x Tgfbr2^f/f^ (Tgfbr2^ΔDC^) BM chimeric mice were inoculated with 1 x 10^6^ live MCA101-OVA cells (s.c.). **E**. Kinetics of tumor growth in Tgfbr2^f/f^ and Tgfbr2^ΔDC^ mice. **F**. Histogram overlay and bar graph depicting H-2K^b^-SIINFEKL staining and mean fluorescence intensity (MFI) on tumor DCs from Tgfbr2^ΔDC^ mice or Tgfbr2^f/f^ controls on day 20 after tumor cell injection. **G**. FACS plots and bar graph showing frequencies of IFN-γ^+^ tumor-infiltrating CD8 T cells in Tgfbr2^ΔDC^ or Tgfbr2^f/f^ animals on day 20 post tumor challenge. **H**. Immunoblot of pSAP and saposins in tumor DCs from Tgfbr2^ΔDC^ or Tgfbr2^f/f^ mice on day 20 after tumor cell inoculation. pSAP-75 = hyperglycosylated prosaposin, pSAP-65 = glycosylated pSAP, and SAPs = saposins. **I**. Differentially expressed glycosyltransferase and glycosidase genes in tumor DCs isolated from Tgfbr2^ΔDC^ or Tgfbr2^f/f^ mice on day 20 after tumor challenge. Splenic DCs from Tgfbr2^f/f^ animals were used as control to calculate mRNA fold change. Data shown in all graphs are representative of three independent experiments, and *p*-values were determined using unpaired Student’s t-test. *p < 0.05; **p < 0.01; ***p < 0.001; ****p < 0.0001.

### Immunotherapeutic targeting of tumor DCs with recombinant prosaposin

Based on the importance of prosaposin for antigen presentation in the tumor microenvironment, we then aimed at targeting recombinant prosaposin to tumor DCs. For this purpose, we coupled prosaposin to anti-DEC205, an antibody well established to target the endocytic receptor DEC205 on DCs, using chemical conjugation as previously described (*22, 23*). Briefly, we incubated recombinant prosaposin with TCEP-HCl (tris (2-carboxyethyl) phosphine hydrochloride) in order to expose its sulfhydryl groups, and in parallel, the anti-DEC205 antibody or isotype control IgG were activated for chemical conjugation by sulfo-SMCC (sulfosuccinimidyl 4-[N-maleimidomethyl] cyclohexane-1-carboxylate). Following overnight incubation, the prosaposin/antibody conjugates were concentrated and successful coupling analyzed by immunoblot (fig. S6A). In addition, we controlled that the amount of prosaposin coupled to anti-DEC205 or isotype control was comparable (fig. S6B). Moreover, we verified that prosaposin conjugation still preserved the fine specificity of the DEC205 antibody by showing that the pSAP/anti-DEC205 conjugate stained a similar percentage of DCs when compared to separate anti-DEC205 detection using flow cytometry (fig. S6C). Furthermore, we incubated pSAP-KO DCs with the prosaposin/antibody conjugates and tested OVA epitope expression and CD8 T cell activation after pulsing of DCs with apoptotic MCA101-OVA cells (fig. S6D). Accordingly, we observed reconstitution of antigen presentation (H-2k^b^-SIINFEKL) and CD8 T cell priming (CD69) specifically by prosaposin coupled to anti-DEC205 in a dose-dependent manner (fig. S6E, F).

After validation of the targeting tool in vitro and ex vivo, we inoculated pSAP-KO BM chimeric mice with MCA101-OVA tumor cells and injected 100 μg prosaposin coupled to anti-DEC205 or isotype IgG on day 9 and 13 after tumor challenge (Fig. 5A). On day 14, we isolated and sorted intra-tumoral DCs, focusing on the two major classical DC subsets cDC1 and cDC2. Staining for intracellular prosaposin in the respective DC subsets revealed effective delivery of pSAP targeted via DEC205 when compared to isotype control (Fig. 5B). This demonstrated successful reconstitution of pSAP-deficient DCs in the tumor microenvironment, especially in the cDC1 subtype that is central to cross-priming of CD8 T cells. Next, we followed a similar experimental schedule using pSAP targeting of tumor-inoculated WT animals (Fig. 5C). Treatment with pSAP coupled to anti-DEC205 strikingly reduced tumor burden in WT mice (Fig. 5D). Correspondingly, antigen presentation of OVA peptide by tumor DCs as well as frequency of IFN-γ-producing, antigen-specific T cells at the tumor site and in draining lymph nodes (dLNs) were significantly amplified upon pSAP treatment (Fig. 5E-G). While pSAP delivery to tumor DCs promoted an immunologically active TME, we next asked whether prosaposin can also rescue immune suppression in immunologically cold tumors. For this purpose, we used the B16F10 melanoma model that displays limited T cell infiltration and low susceptibility to treatment with immune checkpoint inhibitors, such as anti-PD-1, despite strong expression of PD-L1 (*24, 25*). When compared to mice treated with anti-PD-L1 alone, the tumor growth kinetics revealed that pSAP combination therapy was able to overcome the resistance of B16F10 melanoma to immune checkpoint blockade in order to enable protection (Fig. 5H). Taken together, these results highlight the crucial impact of prosaposin on antigen presentation by tumor DCs to trigger powerful intra-tumoral T cell responses and point to a viable future strategy for immunotherapy of cancer.

**Fig. 5.**
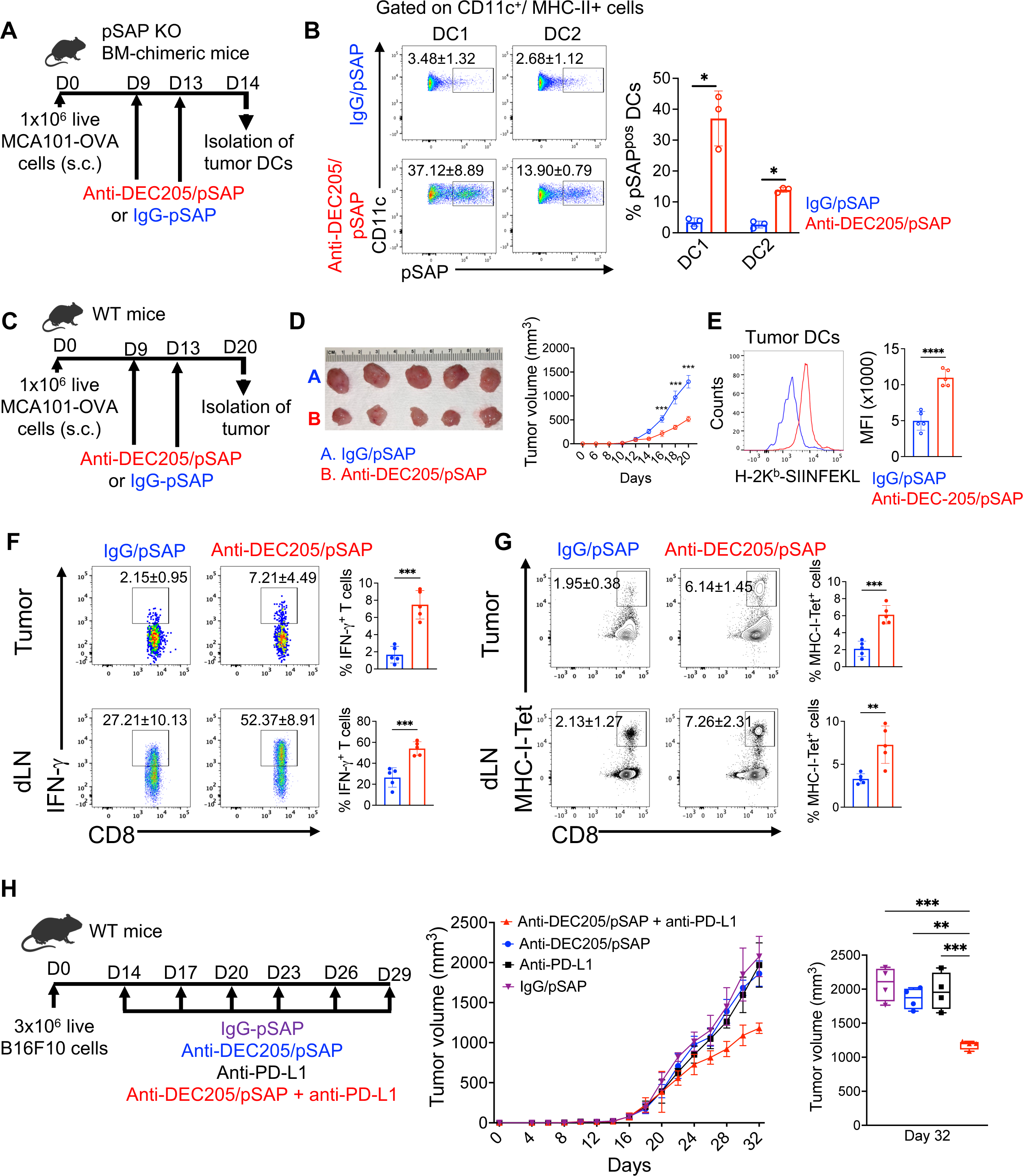
Reconstitution of tumor DCs with recombinant prosaposin drives protection from cancer. **A**. Regime of pSAP targeting to tumor DCs. pSAP KO BM-chimeric mice were inoculated with 1 x 10^6^ live MCA101-OVA cells. On day 9 and 13 after tumor cell injection, mice were intravenously treated with pSAP coupled with either anti-DEC205 or isotype control antibodies. **B**. FACS plots and bar graph showing the amount of prosaposin uptake by DCs at the tumor site as analyzed on day 14 after tumor challenge. **C**. Experimental set-up depicting tumor cell inoculation and pSAP targeting via DEC205 in WT mice. **D**. Comparison of tumor sizes on day 20 (left) and kinetics of tumor growth (right). **E**. Histogram overlay and bar graph showing H-2K^b^-SIINFEKL peptide surface staining and MFI on tumor DCs on day 20 post tumor challenge. **F**. FACS plots and bar graphs showing percentages of IFN-γ-positive CD8 T cells in tumors and tumor-draining lymph nodes (dLN) in mice treated with pSAP coupled to anti-DEC205 or isotype control. **G**. Flow cytometry and bar graphs showing MHC-I (K^b^-SIINFEKL) tetramer-positive CD8 T cells in tumors and draining lymph nodes (dLN). **H**. Experimental set-up depicting B16F10 melanoma cell inoculation, pSAP targeting via DEC205, and tumor growth kinetics. WT mice were inoculated with 3 x 10^6^ live melanoma cells and were treated with pSAP coupled with either anti-DEC205 or isotype control antibodies, either alone or in combination with anti-PD-L1 antibodies. Statistical analysis of tumor volume across all treatment groups is shown on day 32 after tumor challenge. Data shown in all graphs are representative of three independent experiments, and *p*-values were determined using unpaired Student’s t-test. **P < 0.01; ***P < 0.001; ****P < 0.0001.

## Discussion

Herein, we demonstrate the critical function of saposins in processing of corpuscular, membrane associated antigen required for efficient CD8 T cell activation. Notably, saposins were not essential for cross-presentation of soluble antigen, which mechanistically involves rather an early endosomal or phagosomal compartment (*6*). In cancer immunity, the provision of particulate antigen derived from dying tumor cells represents a physiologic and potent route of antigen delivery. In this context, it has been shown that only DCs that contain tumor-derived vesicles are able to induce T cell responses (*26*). Thus, our findings highlight the importance of the lysosome in cross-priming of T cells and underscore the impact of saposins on the immunogenic pathway of vesicular processing.

In tumor biology, previous reports suggested a trophic function of prosaposin able to stimulate proliferation of cancer cells (*27, 28*). However, prosaposin and a pSAP-derived synthetic cyclic peptide were able to prevent tumor metastasis in a mouse model (*29, 30*). Our work established a genuine immunological function of prosaposin in tumor DCs. Thus, pSAP-driven antigen processing and presentation at the tumor site amplified abundance and functionality of tumor infiltrating T lymphocytes, ultimately leading to cancer control.

TGF-β is a pleiotropic cytokine with a diverse set of immunosuppressive functions and is also produced by most cell types. Therefore, tumors themselves as well as the infiltrating cells of the tumor microenvironment can serve as source and target of TGF-β. For example, tumor-derived TGF-β is capable of restricting T cell infiltration or to functionally block differentiation of protective T lymphocyte populations (*15, 31*). Our results revealed that TGF-β acts on tumor DCs to trigger hyperglycosylation of prosaposin and its subsequent secretion, depleting the lysosomal pool of saposins required for proper antigen processing and presentation. The secretion of prosaposin has been shown to be caused by pSAP oligomerization in a cell line (*32*). However, in primary DCs in the tumor microenvironment, we did not observe large oligomers and instead found perturbed interaction between sortilin and hyperglycosylated prosaposin. An additional lysosomal player, called progranulin, might be involved in this mechanism, as it has been shown to bridge the interaction between pSAP and sortilin (*33*). Overall, tumors are prone to immune escape, and our work describes a further cancerous strategy to manipulate an antigen processing molecule via glycosylation.

Since hyperglycosylation of prosaposin occurs along the secretory pathway, an approach to overcome tumor-induced saposin deficiency entails the feeding of recombinant pSAP into the endocytic route of DCs. In this context, our targeting experiments using prosaposin coupled to anti-DEC205 demonstrated that reconstitution with fully functional pSAP is able to restore antigen presentation in tumor DCs, leading to amplified T cell responses and eventual tumor protection. DCs are not only crucial for T cell priming in draining lymph nodes. A growing body of evidence indicates that tumor-associated DCs have vital functions with regard to recruitment of effector T cells and stimulation of tumor-infiltrating T lymphocytes, overall required for effective immunity to cancer (*34–37*). Current therapeutic options, such as immune checkpoint blockade, aim at reinvigorating exhausted T cells to augment anti-tumor responses (*38*). However, these boosted T lymphocytes need to encounter their respective antigens in order to deploy their functions in a tumor-specific manner. Consistent with that notion, we showed that pSAP combination treatment is able to override the resistance of immunologically cold tumors to immune checkpoint inhibitors. Taken together, a prosaposin-based therapy could help to restore powerful antigen presentation by tumor DCs with the goal to drive protective immune responses at the site of the cancer.

## Author contributions

P.S. and F.W. designed the study and performed data analysis. P.S., X.Z., K.L., J.H.K., Q.W., J.K., M.L., and L.K. performed experiments. L.T. produced recombinant prosaposin and saposins. P.S. and F.W. wrote the manuscript.

## Acknowledgements

We thank Dr. Clotilde Théry for providing us with OVA-expressing MCA101 fibrosarcoma cells. We are grateful to Drs. Richard Cummings and Sylvain Lehoux, and all members of the National Center for Functional Glycomics (NCFG) at Harvard Medical School for performing glycan mass spectrometry. This work was supported by National Institutes of Health grants R01 AI136939 to F.W.

## Data and material availability

All data are available in the manuscript or the supplementary material.

## Supplementary material

### Materials and Methods

#### Mice

Wild-type C57BL/6J, B6.SJL (B6.SJL-Ptprca Pepcb/Boy), OVA-specific OT-1 TCR-transgenic mice (B6.TcraTcrb 1100Mjb/J), CD11c-Cre (B6.Cg-Tg(Itgax-cre)1-1Reiz/J), and floxed TGF-β receptor II mice (B6;129-Tgfbr2tm1Karl/J) were purchased from Jackson Laboratories. B6.SJL and OT-I transgenic mice were crossed in our lab to obtain OTI.SJL mice, while CD11c-Cre animals were crossed with floxed TGF-β receptor II (Tgfbr2^f/f^) mice. The generation of prosaposin deficient mice (pSAP-KO) has been reported previously (*39*). All animals were maintained in the animal barrier facility of Harvard Medical School (HMS), and all animal procedures were approved by the IACUC at HMS (approval number IS00001618).

#### Cells

OVA-expressing MCA101 fibrosarcoma cells were developed by Dr. Clotilde Théry (Institut Curie, Paris, France) and have been described previously (*40*). In brief, MCA101/OVAC1C2 expresses the model antigen ovalbumin coupled to the C1C2 domain of milk fat globule EGF factor VIII (MFG-E8)/lactadherin, which targets ovalbumin to PS-expressing vesicles. B16F10 melanoma cells were procured from American Type Culture Collection (USA). Tumor cells were cultured in Dulbecco’s Modified Eagle’s Medium (DMEM) supplemented with 10% FBS, β-mercaptoethanol (55 mM; Gibco), penicillin (100 U/ml) and streptomycin (100 ug/ml), sodium pyruvate (1 mM), HEPES (100 mM), and hygromycin (300 μg/ml; Gibco). DC2.4 cells were purchased from Millipore Sigma and were cultured in complete RPMI medium containing 10% FBS, penicillin (100 U/ml) and streptomycin (100 ug/ml), sodium pyruvate (1 mM), L-glutamine (2 mM), β-mercaptoethanol (50 µM), and HEPES (100 mM). All cells were cultured in an incubator at 37°C, 5% CO_2_, and 95% relative humidity.

#### Antibodies

Antibodies directed against mouse CD8 (53-6.7), CD3 (145-2C11), and Fc block reagent (anti mouse CD16/32) were purchased from BD Biosciences (USA). Antibodies against murine CD45.1 (A20), CD103 (2E7), LAMP-1 (1D4B), CD11b (M1/70), F4/80 (QA17A29), CD24 (QA20A91), IFNγ (XMG1.2), CD11c (N418), IA/IE (M5/114.15.2), XCR1 (ZET), SIRP1α (P84), K^b^-SIINFEKL (25-D1.16) were purchased from BioLegend (USA). Antibodies against human CD146 (P1H12), CD8 (SK1), CD11c (Bu15), CD11b (QA20A58), and LAMP-1 (H4A3) were from BioLegend (USA). Anti mouse DEC205 (NLDC-145) and isotype control (BE0094) antibody were purchased from BioXCell (USA). Anti-mouse and human pSAP antibody (polyclonal) were purchased from Proteintech (Japan), and anti-mouse/human sortilin (polyclonal) was purchased from Abcam (USA). Fluorochrome-labeled HLA-A2 tetramers loaded with peptides from MART (AA 26-35; ELAGIGILTV), NY-ESO-1 (AA 157-165; SLLMWITQV), tyrosinase (AA 369-377; YMDGTMSQV), and gp100 (AA 209-217; ITDQVPFSV) were procured from Tetramer Shop (Denmark).

#### Transmission Electron Microscopy (TEM)

Apoptotic vesicles were generated from the methylcholanthrene (MCA)-induced murine fibrosarcoma cell line MCA101. Cells were washed thoroughly, resuspended in complete RPMI 1640 medium, and subjected to irradiation using a cesium source (100 Gy). Successful apoptosis induction was confirmed by flow cytometry using annexin V staining. Subsequently, vesicles were purified from cell culture supernatants using differential ultracentrifugation to generate 100,000 *g* pellets as previously described (*21*). For electron microscopy, apoptotic bodies were fixed in 4% paraformaldehyde and 2.5% glutaraldehyde and embedded in Spurr’s resin according to the manufacturer’s protocol (Millipore Sigma). Sections were contrasted using lead citrate and analyzed using a Zeiss EM10 microscope.

#### Calcein leakage assay

Apoptotic bodies were first resuspended in a buffer containing 50 mM calcein and subsequently passed through a liposome extruder using a membrane with 100 nm pores. Upon extrusion, the∼400 nm vesicles were forced to transiently open their lipid bilayers to change their size according to the 100 nm pores used. During this process, extruded vesicles incorporated the calcein fluorochrome. To test saposin-induced vesicle disintegration, calcein-loaded apoptotic bodies were incubated with the four different saposins (5 ug/ml) in 50 mM sodium acetate buffer at a pH of 4.5 to facilitate saposin activity. Saposin-induced calcein release corresponded with increased fluorescence, which was measured using a spectrofluorometer. As a negative control, apoptotic bodies were incubated with bovine serum albumin (BSA). Percentage of calcein leakage was calculated using the fluorescence signal as a proportion of maximum calcein release induced by Triton X-100 and the minimum baseline levels.

#### Generation of murine bone marrow-derived dendritic cells

Mouse tibiae and femora were isolated and flushed with ice-cold RPMI 1640 containing 5% FBS, 100 mM HEPES, and cells were then filtered through a 70 µm cell strainer. Red blood cells were lysed using red blood cell lysis buffer (155 mM NH_4_Cl, 10 mM KHCO_3_, 0.1mM EDTA), and bone marrow cells were then plated at a density of 6-8 x 10^6^ cells per 10 cm petri dish in complete RPMI 1640 medium (10% FBS, 100 U/ml penicillin and 100 ug/ml streptomycin, 2 mM L-Glutamine, 1 mM sodium pyruvate, and 0.0012% 2-mercaptoethanol) in the presence of GM-CSF (20 ng/ml, PeproTech). Cells were cultured at 37°C, 5% CO_2_, and 95% relative humidity for 7 days. Thereafter, floating BM-derived dendritic cells (BMDCs) were harvested by gently flushing the cells from the plate, and cell viability was assessed using trypan blue staining.

#### Confocal microscopy

For confocal imaging, MCA101 cells were labeled with 5 μM CFSE and were γ-irradiated to generate apoptotic bodies as described above. 1 x 10^5^ BMDCs were cocultured with 5 x 10^5^ CFSE-labeled apoptotic MCA101 cells for different incubation times in poly-L-lysine-treated 8 chamber slides (Millipore Sigma) at 37°C, 5% CO_2_, and 95% relative humidity. BMDCs were then permeabilized with 0.2% Triton X-100, blocked with PBS containing 2% FBS and 10% goat serum. Cells were further stained overnight with rabbit anti-LAMP1 antibody (Abcam, cat. no.: Ab24170) followed by staining with anti-rabbit-Alexa555 antibody (Molecular Probes, cat. no.: A21429). All samples were mounted with ProLong® Gold antifade mounting medium including DAPI (Life Technologies). Images were taken using an Olympus FV1000 confocal microscope. Images were analyzed using ImageJ/Fiji software v2.0 (NIH). For z-axis image reconstruction, 10 confocal sections, 0.5 μm apart, were assembled using ImageJ/Fiji software.

#### *In vitro* CD8 T cell proliferation assay

Preparation of BMDCs and apoptotic MCA101 cells was performed as described above. Naive CD8 T cells were purified from ovalbumin-specific TCR-transgenic OT-I.SJL mice using a CD8 T cell isolation kit (Miltenyi Biotec). OT-I T cells were suspended in PBS (10^7^ cells/ml) and were incubated with CFSE (Molecular Probes; 5 μM) for 5 min at room temperature, and an equal volume of FBS was added to quench excess CFSE. To test CD8 T cell proliferation, BMDCs were pulsed with apoptotic MCA101 cells at a ratio of 1:5 for 4 hours, and thereafter, 5 x 10^4^ BMDCs were cocultured with 1 x 10^5^ CFSE-labeled OT-I CD8 T cells. In a separate experimental setting, pSAP-deficient BMDCs were pulsed with 10 µg/ml recombinant prosaposin. Cells were cultured in 96-well U-bottom plates at 37°C, 5% CO_2_, and 95% relative humidity for 3 days. The proliferation of CFSE-labeled OT-I CD8 T cells was measured using flow cytometry, while soluble ovalbumin (50 µg/ml) was used as a positive control.

#### Tumor cell inoculation

Mice were subcutaneously injected with 1 x 10^6^ live MCA101 tumor cells in the shaved right flank. In separate experiments, mice were immunized with 4 x 10^5^ irradiated MCA101 tumor cells in the left flank 7 days before live tumor challenge. For both experimental settings, tumor growth was monitored with calipers every 2 days, and a tumor volume of 2,000 mm^3^ was considered as an endpoint.

#### Flow cytometry and cell purification

Subcutaneous tumors were cut into small pieces and were digested in 20 ml of RPMI containing collagenase D (400 μg/ml), collagenase VIII (400 μg/ml), and 2% FCS for 1 hour at 37°C. Subsequently, 70% and 30% percoll density gradients were used to remove debris and dead cells, and single cell suspensions were collected from the gradient interface. Spleens and tumor draining lymph nodes (dLNs) were cut and dissociated with 10 U/ml collagenase I and 30 U/mL DNase I (Millipore Sigma) for 45 min at 37°C, and single cell suspensions were collected by passing cells through 70-μm cell strainers. Red blood cells from all samples were removed using erythrocyte lysis buffer.

For flow cytometry experiments, single cell suspensions from tumor, spleen, and dLN were blocked with mouse Fc block reagent (BD Biosciences) for 15 min at 4°C, and were then stained with antibodies directed against CD45, CD8, LAMP-1, CD11c, CD24, CD103, CD11b, MHC-II, H-2K^b^-SIINFEKL, F4/80, XCR-1, and SIRP1α. For IFN-γ staining, cells were stimulated with PMA (50 ng/ml) and Ionomycin (500 ng/ml; Sigma-Aldrich) for 4 hours in the presence of GolgiStop (BD Biosciences). Next, cells were surface stained, fixed, and incubated with anti-IFN-γ antibody using the Cytofix/Cytoperm kit (BD Biosciences) following the manufacturer’s protocol. For tetramer staining, 2 x 10^6^ cells suspended in 50 µL PBS were incubated with antigen-specific MHC-I OVA tetramer (SIINFEKL; 1:50 dilution) for 15 min at 37°C in the dark. Single cell suspensions from human melanoma samples were surface stained with antibodies against CD146, CD8, CD11b, CD11c, CD45, and LAMP-1, while intracellular staining for IFN-γ was performed using the Cytofix/Cytoperm kit (BD) following the manufacturer’s protocol. In a separate set of experiments, human melanoma samples were stained with fluorochrome-labeled HLA-A2 tetramers loaded with epitopes from MART (AA 26-35; ELAGIGILTV), NY-ESO-1 (AA 157-165; SLLMWITQV), tyrosinase (AA 369-377; YMDGTMSQV), and gp100 (AA 209-217; ITDQVPFSV), incubating 1 x 10^5^ cells with the respective tetramers for 45 min at 37°C in the dark. Data were acquired using a BD FACS Canto II (BD Biosciences) and analyzed with FlowJo software (Treestar).

Cell purifications were performed either by FACS-assisted cell sorting or by magnetic-activated cell sorting (MACS). For magnetic cell separation, cells were suspended in MACS buffer, and DCs and naive CD8 T cells were isolated using the respective beads following the manufacturer’s instructions (Miltenyi Biotec). Human CD146^+^ melanoma cells, CD8^+^ T cells, and CD11c/b^+^ myeloid cells, as well as mouse cDC1 and cDC2 from tumor and spleen were FACS-sorted using a BD FACS Aria II, and purity of cells was determined by post-sort analysis based on flow cytometry.

#### Antigen uptake and processing assays

To test their phagocytosis and antigen processing capacities, DCs isolated from tumor, spleen, or dLNs were incubated with either FITC-dextran (1 mg/ml), or DQ-ovalbumin (0.5 mg/ml) for 1 hour in a CO_2_ incubator at 37°C. DQ-OVA is a self-quenching reagent, which fluoresces only after proteolytic cleavage by lysosomal enzymes. While FITC-dextran uptake defines the extent of endocytosis, any fluorescence in the green channel derived from cleaved DQ-OVA represents the antigen processing abilities of DCs. Background signal was determined by culturing control DCs at 4°C, and mean fluorescence intensity (MFI) was subsequently measured by flow cytometry.

#### *In vivo* CD8 T cell priming

OT-I CD8 T cells were isolated from OT-I.SJL mice using MACS sorting (Miltenyi Biotec) and labeled with 5 μM CFSE in PBS. CFSE-labeled cells were tested for viability using trypan blue staining, and 5 x 10^6^ viable OT-I CD8 T cells were then intravenously administered to WT or pSAP-KO mice. On the following day, 5 x 10^6^ irradiated MCA101-OVA cells were subcutaneously injected into each recipient’s left flank, and draining inguinal lymph nodes (dLN) were harvested four days later. Proliferation of OT-I CD8 T cells and the production of IFN-γ were measured using flow cytometry.

#### Bone marrow chimera generation

Bone marrow (BM) cells were prepared from tibias and femurs of WT, pSAP-KO, CD11c-Cre x Tgfbr2^f/f^ (Tgfbr2^ΔDC^), and littermate control (Tgfbr2^f/f^) mice as described above. WT mice were irradiated using a cesium source (900 rad), and irradiated mice were injected with 1-3 x 10^6^ BM cells from pSAP-KO, Tgfbr2^ΔDC^, or WT littermate control mice. Chimera were treated with 2 mg/ml neomycin (Sigma-Aldrich) in drinking water for 4 weeks. Eight to twelve weeks after BM transfer, mice were assessed for chimerism and subsequently used for tumor experiments.

#### Immunoprecipitation and western blot

FACS-sorted DCs were lysed in 50 mM Tris (pH 8.0), containing 150 mM NaCl, 1% Triton X-100, 0.1% deoxycholic acid and 1x protease inhibitors (Roche). Anti-pSAP antibody was coupled with amine-reactive resin following the manufacturer’s instructions (Thermo Fisher Scientific). Subsequently, cell lysates were incubated overnight with resin-conjugated anti-pSAP antibody, and bound protein complexes were enriched using column purification (Thermo Fisher Scientific) following the manufacturer’s protocol. For immunoblot analysis, cells were lysed in RIPA buffer containing 1x protease and phosphatase inhibitors, and protein concentrations were determined using a BCA protein quantification kit (Bio-Rad). Samples were diluted in Laemmli’s sample buffer (Bio-Rad) and were loaded onto 15% Mini-PROTEAN TGX precast gels (Bio-Rad). Electrophoresis was carried out at 120 V for 1 hour in tris-glycine buffer. Separated proteins were transferred to nitrocellulose membranes, and the membranes were blocked in 5% BSA dissolved in TBST buffer (tris-buffered saline, containing 20 mM Tris-HCl, pH 7.6, 150 mM NaCl, 0.1% Tween 20). After blocking, membranes were incubated with antibodies against either mouse pSAP, sortilin, β−actin, or human pSAP at 4°C overnight. Thereafter, membranes were washed three times with TBST and were incubated with 1:5,000 diluted HRP-labeled anti-rabbit antibody (Thermo Fisher Scientific) for 1 hour. After additional three washes, membranes were exposed to chemiluminescent substrate and imaged upon exposure to X-ray films in the dark.

#### Endoglycosidase H (Endo H) digestion assay

FACS-sorted DCs were lysed in RIPA buffer, and cell lysates were subjected to Endo H digestion according to the manufacturer’s protocol (New England Biolabs). Briefly, 100 μg of protein lysate was denatured using the glycoprotein-denaturing buffer supplied by the manufacturer for 10 minutes at 99°C. Subsequently, samples were cooled at 4°C, and 500 U of Endo H was added to each sample. Glycan digestion was carried out at 37°C for at least 12 hours, and the release of glycans from pSAP was determined by immunoblot analysis.

#### ELISA

pSAP concentrations were determined using sandwich ELISA following the manufacturer’s instructions (Abcam). Briefly, plates precoated with antibody specific for pSAP were incubated with standards and samples for 2 hours at room temperature. The wells were washed three times, and biotinylated anti-mouse pSAP antibody was added to each well (1:500 dilution). Thereafter, plates were washed three times, and HRP-conjugated streptavidin was added to each well, prior to incubation for 1 hour at room temperature. TMB substrate solution was added to each well, and the color reaction was stopped by using 2N H_2_SO_4_. Plates were then measured for optical density at 450 nm using a spectrophotometer.

#### Proximity ligation assay (PLA)

FACS-sorted DC subsets from tumor and spleen were plated on eight-chambered slides for 45 min at 37°C, and adherent cells were washed and fixed for 15 min with 2% paraformaldehyde. Cells were permeabilized with 0.5% saponin in PBS for 30 min. PLA was carried out with Duolink® *in situ* detection reagents following the manufacturer’s instructions (Millipore Sigma). Anti-pSAP antibody was labeled with plus probes, while anti-sortilin antibodies were labelled with minus probes. For PLA reaction, cells were blocked with Duolink® blocking solution for 1 hour at room temperature and subsequently incubated with the respective PLA probes for 2 hours. After washing cells three times, all probe-labeled cells were subjected to PLA signal amplification. Finally, cells were mounted on slides with DAPI-containing mounting reagent. Images were taken using an Olympus Fluoview (FV) 1000 confocal microscope, and data were analyzed using ImageJ software (NIH).

#### qRT-PCR for glycosyltransferase and glycosidase genes

CD11c^+^ DCs from spleen and tumor were purified using FACS sorting. DC2.4 cells were collected upon trypsinization after 4 days of 2.5 ng/ml TGF-β stimulation. Expression of genes involved in glycosylation were quantified by quantitative real-time PCR (qRT-PCR) using the glycosylation RT^2^ Profiler PCR Array according to the manufacturer’s instructions (PAMM-046Z, Qiagen). Briefly, RNA was isolated from DCs using the RNeasy Kit (Qiagen), and 50 ng RNA was converted to cDNA using the RT^2^ First Strand Kit (Qiagen). The cDNAs were subjected to amplification using gene-specific primers and SYBR Green (RT^2^ SYBR Green Master Mix, Qiagen) in an Ab7100 Real-Time PCR System (Thermo Fisher Scientific). Amplification of the gene encoding for β-actin served as internal control, and relative gene expression levels were calculated using the 2^-ΔΔCT^ method.

#### Mass Spectrometry

FACS-sorted CD11c^+^ DCs from spleen and tumor were lysed, and pSAP was immunoprecipitated as described above, prior to loading onto 12% acrylamide gels and visualization using coomassie blue staining. pSAP bands corresponding to a molecular weight of 65 kDa and 75 kDa were excised and incubated at room temperature for 5 min with 200 µl of a freshly prepared 50 mM (3.91 mg/ml) ammonium bicarbonate solution and 200 µl of acetonitrile (100%). Subsequently, samples were dried using a SpeedVac and incubated with freshly prepared 10 mM (1.54 mg/ml) 1,4-dithiothreitol (DTT, Sigma) at 50°C for 30 min. Thereafter, pSAP was alkylated with 55 mM (10.2 mg/ml) iodoacetamide (IAA, Sigma, #I6125) at room temperature for 30 mins, and peptides were subsequently cleaved by incubating samples with a 20 µg/ml solution of TPCK-treated trypsin (Sigma, #4352157) at 37°C overnight. Trypsin digestion was stopped by the addition of ∼2 drops of 5% acetic acid, and samples were added to a C18 Sep-Pak (200 mg) column (Waters) preconditioned with a solution of methanol, 1-propanol, and 5% acetic acid (2:2:1). Reaction tubes were washed with 1 mL of 5% acetic acid and added to the column, followed by an additional 5 mL wash of 5% acetic acid. Each column was placed in a 15 mL glass tube, and glycopeptides were eluted using 2 mL of 20% 1-propanol, 2 mL of 40% 1-propanol, and 2 mL of 100% 1 propanol. The eluted fractions were pooled and placed in a SpeedVac to remove the excess organic solvent, followed by lyophilization. For N-glycan release, dried peptides were treated with 1 ml of PNGase F in 200 µl of 50 mM ammonium bicarbonate solution overnight at 37°C. Permethylated glycans were resuspended in 25 µL of 75% methanol and spotted in a 1:1 ratio with DHB matrix on an MTP 384 polished steel target plate (Bruker Daltonics) as previously described (*41*). MALDI-TOF MS data were acquired based on a Bruker Ultraflex II instrument using Flex Control software in the reflective positive mode. For N-glycans, a mass/charge (m/z) range of 1,000 - 5,000 kDa was collected, and twenty independent captures were obtained from each sample and averaged to create the final combined spectra file. Data were exported in .msd format using Flex Analysis software for subsequent annotation.

#### TGF-β in vitro stimulation

DC2.4 cells were seeded in complete RPMI at 75% confluency in 24-well plates. The following day, cells were washed and cultured in complete RPMI containing 0, 2.5, or 10 ng/ml recombinant TGF-β for 4 days. Culture supernatant was concentrated using 10 kDa protein concentrators, and the amount of prosaposin released was quantified using ELISA.

#### Melanoma Patient Samples

Dissociated whole tumor samples were purchased from Discovery Life Sciences (USA). Melanoma, myeloid, and CD8 T cells were FACS-sorted as described above. Melanoma cells were γ-irradiated (10,000 rad), and induction of apoptosis was monitored by Annexin V staining using flow cytometry. 30,000 DCs were cocultured with 150,000 CD8 T cells (1:5 ratio), and 1 x 10^5^ irradiated melanoma cells were used as antigen source. To test the function of pSAP, cultures were stimulated with 5 μg recombinant human pSAP (Abcam). Cells were cocultured for 5 days, and CD8 T cell activation was quantified by cytokine and tetramer staining using flow cytometry as described above.

#### Generation of monocyte-derived DCs

Peripheral blood mononuclear cell samples were procured from Discovery Life Sciences (USA). Monocytes were isolated from mononuclear cell fractions using CD14 magnetic microbeads, according to the manufacturer’s instructions (Miltenyi Biotec). Isolated monocytes were cultured in tissue-culture dishes at 0.4 x 10^6^ cells/mL in complete RPMI 1640 medium, containing recombinant human IL-4 (50 ng/mL; R&D Systems) and recombinant human GM-CSF (100 ng/mL; R&D Systems), with 1 supplement of fresh medium and cytokines on day 3 of culture. Cells were harvested on day 6 to be subsequently used for immunoblotting and PLA.

#### Chemical coupling of pSAP with anti-DEC205 antibody

Recombinant pSAP was coupled with anti-DEC205 antibody activated with sulfo-SMCC (sulfosuccinimidyl 4-(N-maleimidomethyl) cyclohexane-1-carboxylate; Pierce Chemical Co.), following the manufacturer’s protocol. pSAP was reduced with TCEP (Tris[2-carboxyethyl] phosphine hydrochloride) immobilized onto 4% crosslinked beaded agarose for 90 min at room temperature. In parallel, anti-DEC205 or isotype control antibodies were maleimide-activated by incubation with sulfo-SMCC for 30 min at 37°C. Excess of salts were removed by desalting columns, and subsequently, reduced pSAP and maleimide-activated antibodies were incubated overnight at 4°C. Antibody/pSAP conjugates were purified by passing through 150 kD protein concentrators (Thermo Fisher Scientific), and the purity and concentrations of antibody/pSAP conjugates were determined using SDS-PAGE and ELISA. To test binding of anti-DEC205/pSAP to the DEC205 receptor, pSAP-KO BMDCs were incubated with anti-DEC205/pSAP conjugates, prior to staining with anti-pSAP antibody. Isotype antibody-coupled pSAP was used as negative control, while direct staining of DCs with anti-DEC205 antibody was used as positive control. To examine CD8 T cell activation, pSAP-KO BMDCs were pulsed with irradiated MCA101-OVA cells and cocultured with OT-I CD8 T cells in the presence of either anti-DEC205/pSAP or isotype/pSAP conjugates. To investigate the in vivo delivery of anti-DEC205/pSAP to tumor DCs, MCA101 tumor-bearing pSAP-KO BM chimeric mice were injected with 100 μg pSAP conjugated to either anti-DEC205 or isotype antibodies on days 9 and 13 after tumor cell inoculation. pSAP delivery to cDC1 and cDC2 was quantified using flow cytometry. In another set of experiments, B16F10-harboring mice were injected with 100 μg anti-DEC205/pSAP conjugate combined with 100 μg of either anti-PD-L1 or isotype control antibody for six injections every three days.

#### Statistical analyses

All data are presented as mean and standard deviation of the mean (SD). The statistical significance of differences was calculated using a two-tailed Student’s t-test for both paired and unpaired samples using Prism 9.0 (GraphPad software). p-values ≤ 0.05 were considered significant (\**p* < 0.05; \*\**p* < 0.01; \*\*\**p* < 0.001; \*\*\*\**p* <0.0001).

## >Supplementary Figure Legends

**Fig. S1.**
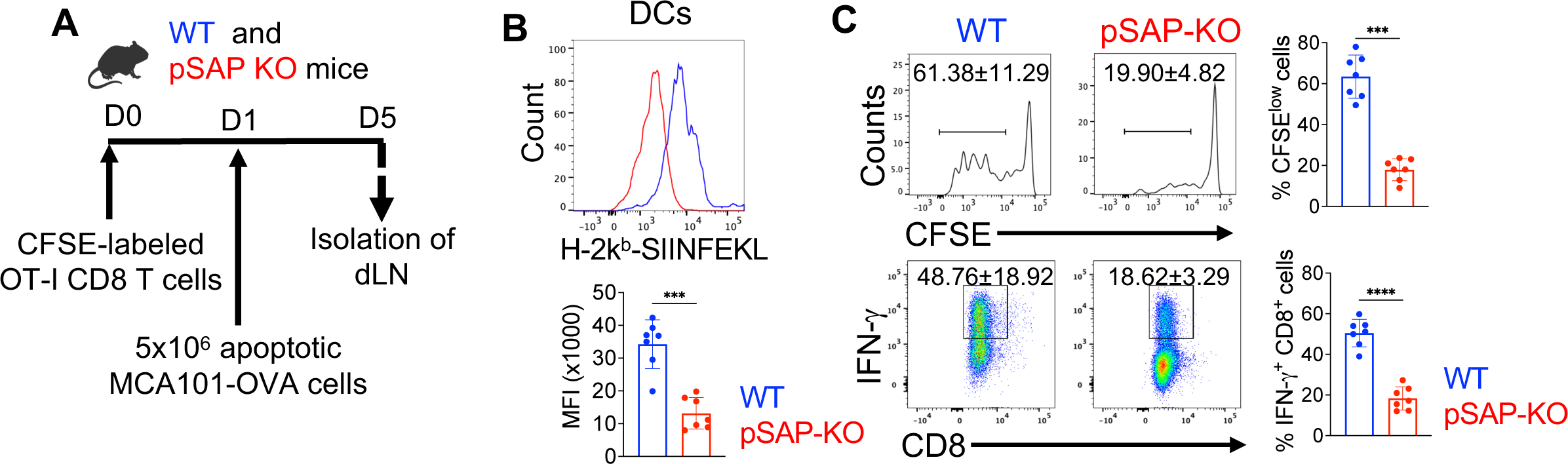
Prosaposin is required for cross-priming of CD8 T cells. **A**. Experimental set-up. WT and pSAP-KO mice were intravenously injected with CFSE-labeled OT-I T cells, followed by subcutaneous administration of 5 x 10^6^ irradiated MCA101-OVA cells on day 1 after adoptive T cell transfer. iLN = inguinal lymph node. **B**. Histogram overlay and bar graph showing H-2K^b^ SIINFEKL peptide staining and mean fluorescence intensity (MFI) on migratory DCs. Migratory DCs were gated as CD11c^+^ and MHC-II^high^ cells. **C**. FACS plots and bar graphs showing frequencies of CFSE^low^ CD8 T cells (top panel) and IFN-γ-positive CD8 T cells (bottom panel) in iLN on day 5 after T cell transfer. Data shown are representative of 7 independent biological replicates, and *p*-values were determined using unpaired Student’s t-test. ***p < 0.001.

**Fig. S2.**
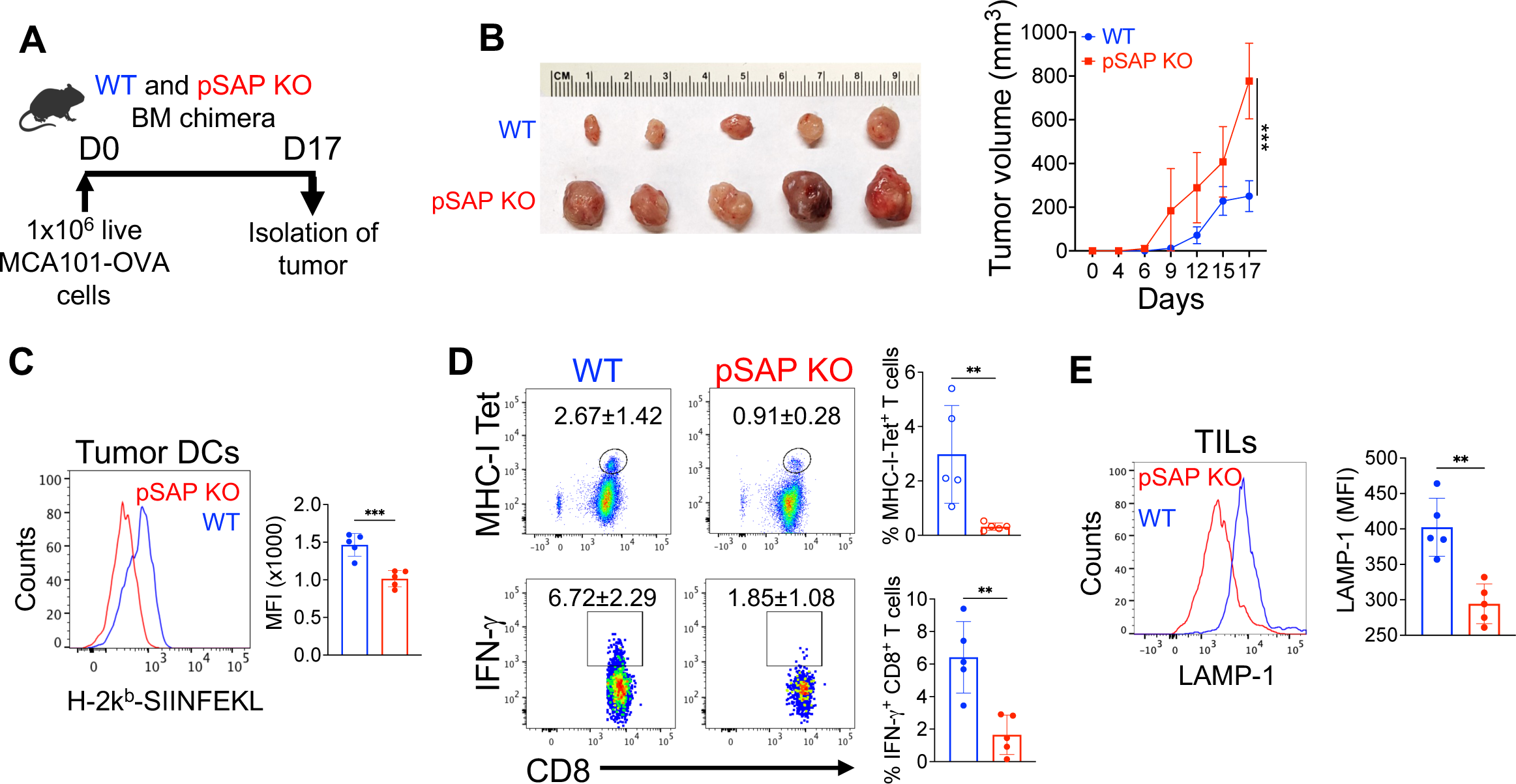
Cancer growth and immunity are controlled by prosaposin. **A.** Experimental scheme of tumor cell injections. WT and pSAP-KO BM chimeric mice were inoculated with 1 x 10^6^ live MCA101-OVA cells (s.c.), and mice were sacrificed 17 days later for tumor harvest and cellular analysis. **B**. Comparison of tumor sizes on day 17 after challenge (left) and kinetics of tumor growth (right). **C**. Histogram overlay and bar graph depicting H-2K^b^-SIINFEKL peptide staining and mean fluorescence intensity (MFI) on tumor DCs. **D**. FACS plots and bar graphs showing frequencies of MHC-I (K^b^-SIINFEKL) tetramer- and IFN-γ-positive tumor-infiltrating CD8 T cells. **E**. Histogram overlay and bar graph depicting LAMP-1 staining and MFI on the surface of tumor infiltrating CD8 T cells. Data shown are representative of three independent experiments. Statistical analysis was performed using unpaired Student’s t-test in graphs C-E. **p < 0.01; ***p < 0.001.

**Fig. S3.**
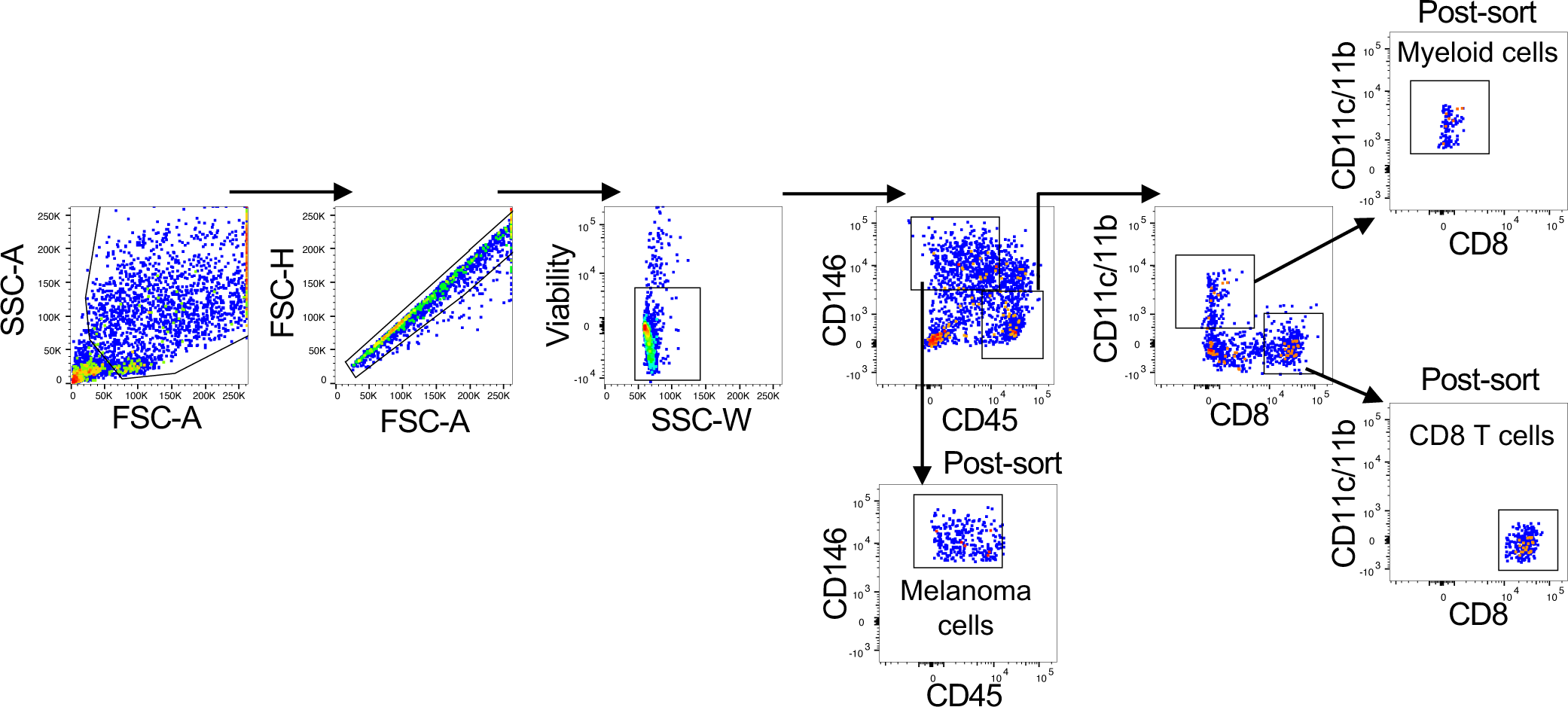
FACS sorting of tumor, myeloid, and T cells from human melanoma patients. FACS plots show the hierarchical gating strategy to identify melanoma (CD146^+^), myeloid (CD45^+^, CD11c/b^+^), and cytotoxic T cells (CD45^+^, CD8^+^), sub-gated from viable cells after doublet discrimination using FSC-A vs. FSC-H gating. Three-way sorting was used to separate the indicated populations, and purity was assessed using post-sort analysis based on flow cytometry.

**Fig. S4.**
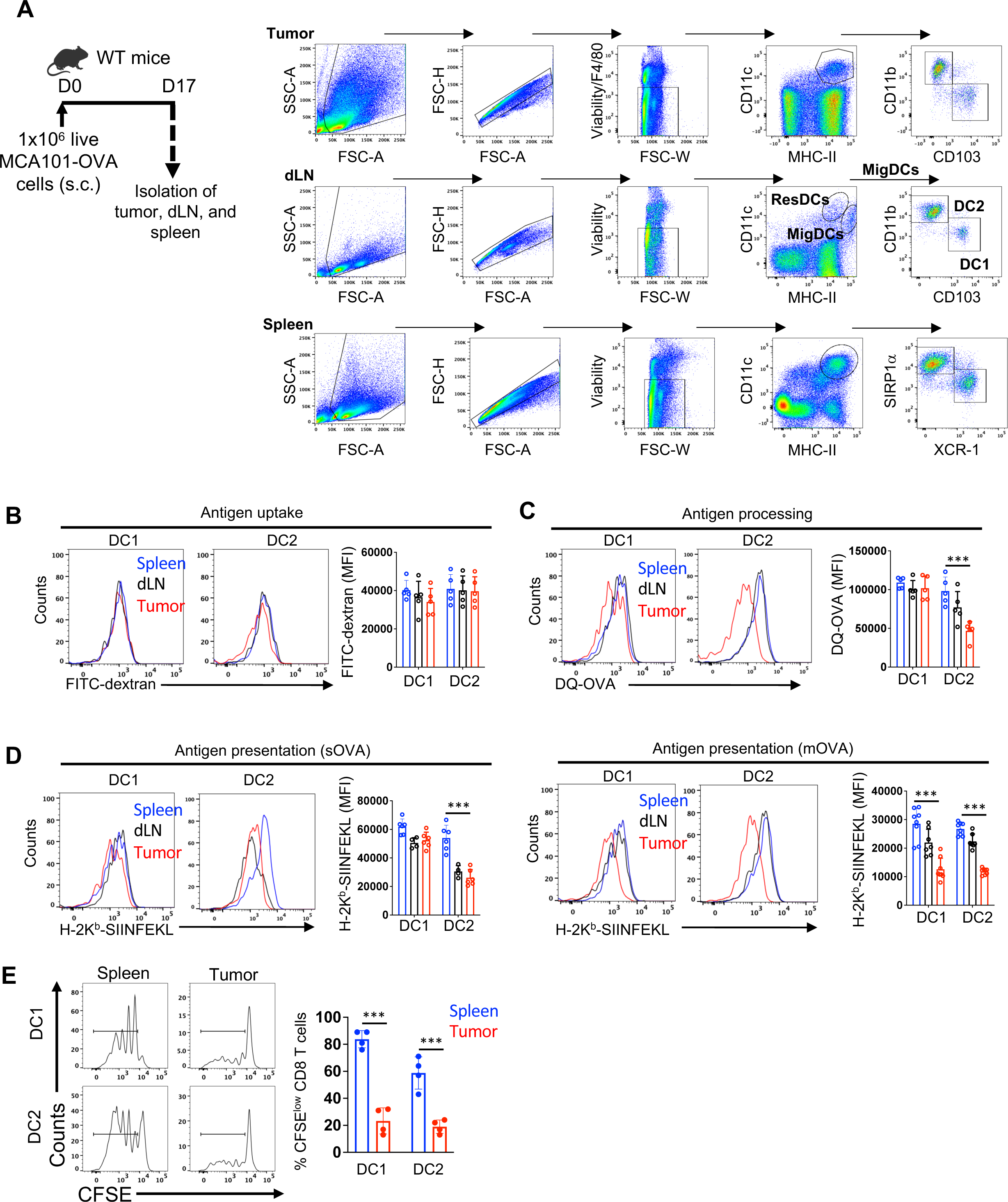
Tumor DCs are impaired in antigen processing and presentation. **A.** Experimental set-up and flow cytometry gating strategy to identify and isolate cDC1 and cDC2 populations from tumor, draining lymph nodes (dLN), and spleen. WT mice were inoculated with 1 x 10^6^ live MCA101-OVA cells (s.c.), and tumor, dLN, and spleen were isolated on day 17 after tumor challenge. ResDCs = resident DCs, MigDCs = migratory DCs. **B**. Antigen uptake assay using FITC-dextran. FACS-sorted DC subsets were incubated with FITC-dextran for 45 minutes, and background signal was quenched by trypan blue staining. Mean fluorescence intensity (MFI) of FITC was quantified using flow cytometry. **C**. Antigen processing assay using DQ-OVA. FACS-sorted DC subsets were incubated with DQ-OVA for 1 hour, and MFI of DQ-OVA was quantified using flow cytometry. DQ-OVA is a self-quenching reagent which fluoresces after proteolytic cleavage in lysosomes. **D**. Antigen presentation assay using soluble OVA or irradiated tumor cells. Purified DCs were pulsed either with soluble OVA (sOVA, left panel), or with γ-irradiated MCA101-OVA cells (mOVA, right panel) for 4 hours. Antigen presentation was quantified using H-2K^b^-SIINFEKL staining on DCs and is depicted in histograms and bar graphs plotting MFI. **E**. T cell assay using OT-I T cells stimulated by cDC1 and cDC2 purified from spleen or the tumor microenvironment (TME). DCs were pulsed with γ-irradiated MCA101-OVA cells for 4 hours and were subsequently cocultured with CFSE-labeled OT-I T cells for 3 days. CFSE dilutions were measured by flow cytometry to assess CD8 T cell proliferation. Data in all graphs show mean±SD, and *p*-values were calculated using one-way ANOVA. ***p < 0.001.

**Fig. S5.**
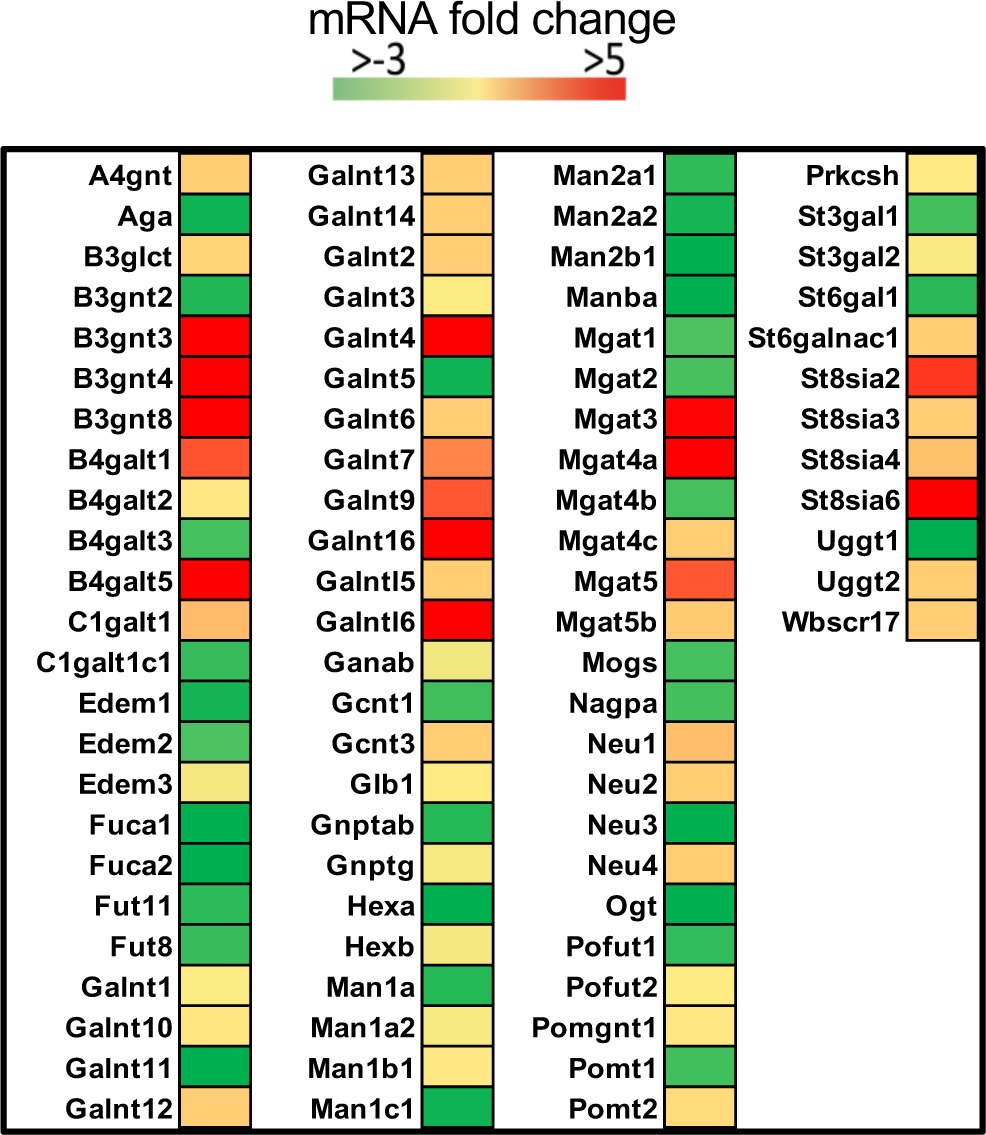
TGF-β treatment changes expression profile of glycosylation genes in DC2.4 cells. Heat map of differentially expressed genes involved in glycosylation in DC2.4 cells after TGF-β treatment. DC2.4 cells were treated with 10 ng/ml TGF-β for 48 hours, and mRNA fold change was determined using the real-time RT^2^ profiler PCR array compared with untreated DC2.4 cells.

**Fig. S6.**
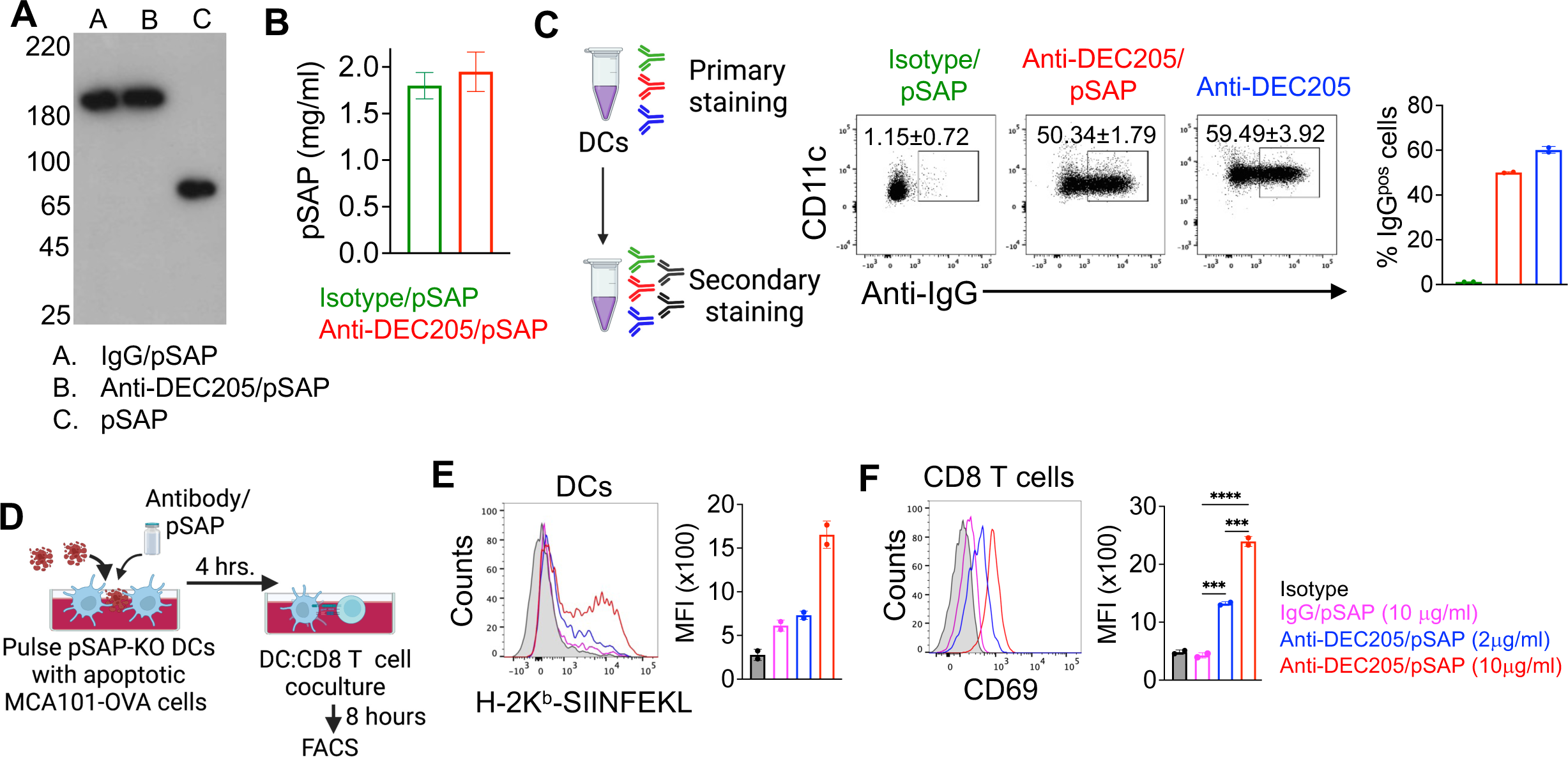
Targeting of pSAP to DCs via anti-DEC205 promotes antigen cross-presentation. **A.** Native electrophoresis and immunoblot of pSAP coupled to isotype or anti-DEC205. Antibody coupled pSAP was detected at molecular weight > 180 kD (lane A and B), while pSAP was detected as 65 kD band (lane C). **B**. ELISA-based quantification of the amount of prosaposin coupled to anti-DEC205 or isotype control antibody. ELISA plates were coated with anti-IgG capture antibodies, followed by incubation with isotype- or anti-DEC205-coupled pSAP. Anti pSAP antibody was used for detection. **C**. FACS plots and bar graph showing engagement of anti-DEC205/pSAP conjugates on the DC surface. In a primary step, BMDCs were incubated for 15 minutes using either isotype- or anti-DEC205-coupled pSAP, or unconjugated DEC205 antibody that served as positive control. In a subsequent step, fluorochrome-tagged secondary antibodies were used to detect pSAP/antibody conjugates or DEC205 on the surface of DCs. **D**. Experimental set-up showing pulsing of pSAP-KO BMDCs with γ-irradiated MCA101-OVA cells, followed by coculture with OT-I T cells in the presence of either isotype- or anti-DEC205-coupled pSAP. **E**. Histogram overlay and bar graph depicting MHC-I-restricted antigen presentation (H-2K^b^-SIINFEKL staining) by pSAP-KO BMDCs pulsed with γ-irradiated MCA101-OVA cells and incubated with either isotype- or anti-DEC205-coupled pSAP using indicated concentrations. Analysis by flow cytometry and quantification using mean fluorescence intensity (MFI). **F**. FACS histogram overlay and bar graph showing CD69 staining and MFI on CD8 T cells cocultured with pSAP-KO BMDCs treated with pSAP/antibody conjugates at indicated concentrations. Data in all graphs depict mean ±SD, and *p*-values were determined using unpaired Student’s t-test. ***p < 0.001; ****p < 0.0001.

**Fig. S7.**
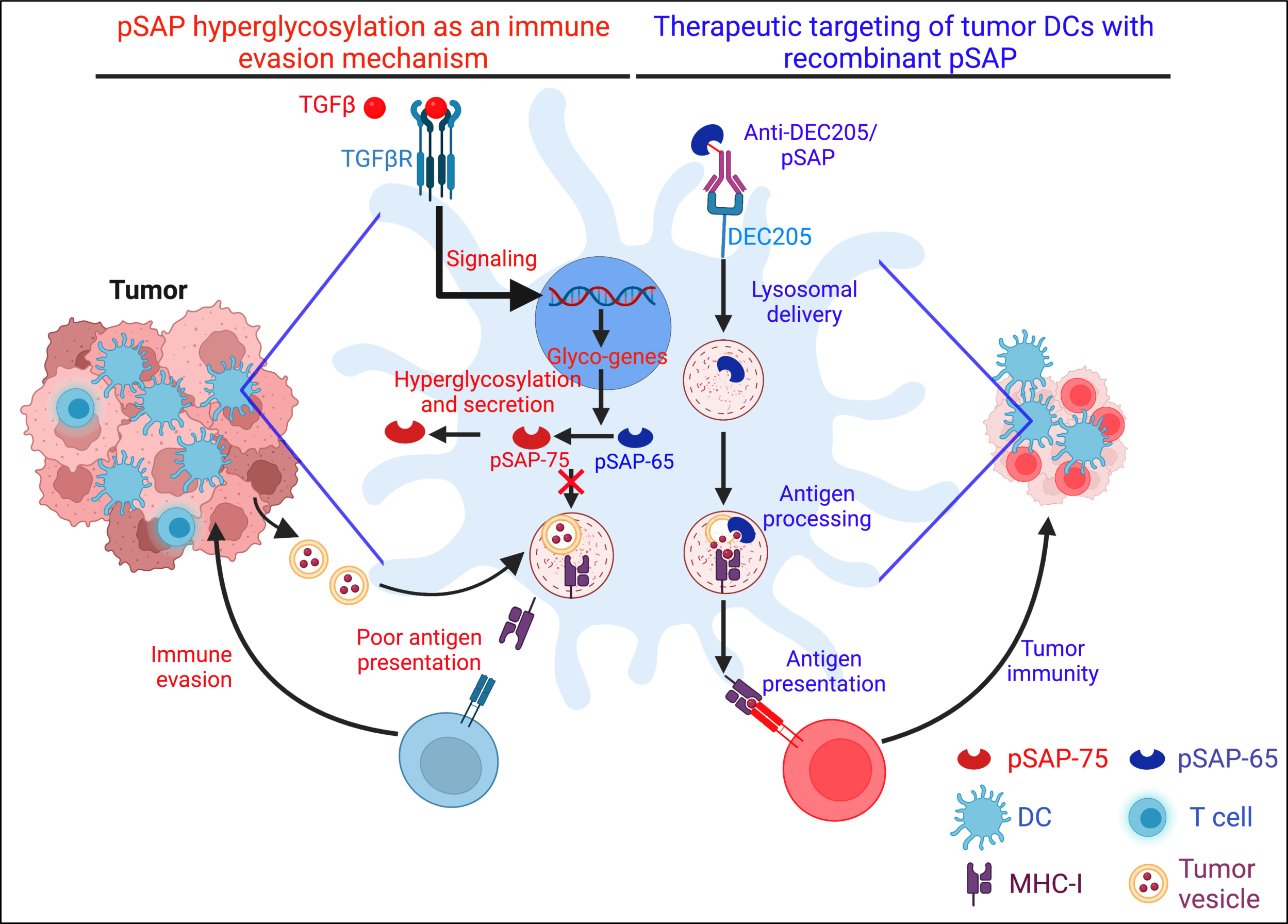
Mechanism of prosaposin-based immune escape and its targeting for immunotherapy of cancer. *Left side*: Hyperglycosylation of prosaposin and immune evasion. TGF-β induces hyperglycosylation of prosaposin in tumor DCs, leading to its secretion. As a consequence, reduced availability of saposins in the endolysosomal compartment impairs antigen processing and presentation, causing hampered T cell responses and loss of tumor control. *Right side*: Therapeutic targeting of tumor DCs with recombinant prosaposin. To overcome immune escape in cancer, recombinant prosaposin can be targeted to endolysosomes in tumor DCs via the endocytic receptor DEC205. Delivery of functional prosaposin restores antigen processing and presentation, triggering reinvigorated T cell responses to promote tumor immunity.

**Table S1.** Description of melanoma patient samples used in this study.

| **Primary diagnosis** | **Tumor location** | **Overall clinical stage** | **Overall treatment status** | **Patient age at collection** | **Gender** | **HLA-A02 by flow cytometry** |
| --- | --- | --- | --- | --- | --- | --- |
| Melanoma | Primary tissue from back skin | III-A | Pre Tx | 67 | Male | Positive |
| Melanoma | Primary tissue from back skin | III-C | Pre Tx | 72 | Female | Positive |
| Melanoma | Primary tissue from neck skin | III | Pre Tx | 61 | Female | Negative |
| Melanoma | Primary tissue from back skin | III | Pre Tx | 69 | Female | Negative |
| Melanoma | Primary tissue from skin | III | Post Tx | 70 | Male | Negative |
| Melanoma | Primary tissue from left hip | III-C | Pre Tx | 68 | Female | Negative |
| Melanoma | Primary tissue from skin | III-C | Pre Tx | 64 | Female | Positive |
| Melanoma | Primary tissue from left arm skin | III | Pre Tx | 57 | Female | Positive |
| Melanoma | Primary tissue from left hip | II-B | Pre Tx | 57 | Male | Positive |
| Melanoma | Primary tissue from neck skin | III-C | Post Tx | 67 | Male | Positive |
| Melanoma | Primary tissue from back skin | III | Pre Tx | 62 | Female | Positive |
| Melanoma | Primary tissue from neck skin | III | Pre Tx | 65 | Female | Positive |

## References

1. M. D. Vesely, M. H. Kershaw, R. D. Schreiber, M. J. Smyth, Natural innate and adaptive immunity to cancer. Annu Rev Immunol 29, 235–271 (2011).

2. M. S. Rooney, S. A. Shukla, C. J. Wu, G. Getz, N. Hacohen, Molecular and genetic properties of tumors associated with local immune cytolytic activity. Cell 160, 48–61 (2015).

3. M. Sade-Feldman et al., Resistance to checkpoint blockade therapy through inactivation of antigen presentation. Nat Commun 8, 1136 (2017).

4. A. Alloatti et al., Critical role for Sec22b-dependent antigen cross-presentation in antitumor immunity. J Exp Med 214, 2231–2241 (2017).

5. J. S. Blum, P. A. Wearsch, P. Cresswell, Pathways of antigen processing. Annu Rev Immunol 31, 443–473 (2013).

6. J. M. Blander, Regulation of the Cell Biology of Antigen Cross-Presentation. Annu Rev Immunol 36, 717–753 (2018).

7. L. Shen, L. J. Sigal, M. Boes, K. L. Rock, Important role of cathepsin S in generating peptides for TAP-independent MHC class I crosspresentation in vivo. Immunity 21, 155–165 (2004).

8. L. Saveanu et al., IRAP identifies an endosomal compartment required for MHC class I cross-presentation. Science 325, 213–217 (2009).

9. T. L. Tang-Huau et al., Human in vivo-generated monocyte-derived dendritic cells and macrophages cross-present antigens through a vacuolar pathway. Nat Commun 9, 2570 (2018).

10. K. Hildner et al., Batf3 deficiency reveals a critical role for CD8alpha+ dendritic cells in cytotoxic T cell immunity. Science 322, 1097–1100 (2008).

11. S. T. Ferris et al., cDC1 prime and are licensed by CD4(+) T cells to induce anti-tumour immunity. Nature 584, 624–629 (2020).

12. T. F. Gajewski, H. Schreiber, Y. X. Fu, Innate and adaptive immune cells in the tumor microenvironment. Nat Immunol 14, 1014–1022 (2013).

13. S. K. Wculek et al., Dendritic cells in cancer immunology and immunotherapy. Nat Rev Immunol 20, 7–24 (2020).

14. R. A. Flavell, S. Sanjabi, S. H. Wrzesinski, P. Licona-Limon, The polarization of immune cells in the tumour environment by TGFbeta. Nat Rev Immunol 10, 554–567 (2010).

15. S. Mariathasan et al., TGFbeta attenuates tumour response to PD-L1 blockade by contributing to exclusion of T cells. Nature 554, 544–548 (2018).

16. S. Lefrancois, J. Zeng, A. J. Hassan, M. Canuel, C. R. Morales, The lysosomal trafficking of sphingolipid activator proteins (SAPs) is mediated by sortilin. EMBO J 22, 6430–6437 (2003).

17. L. Yuan, C. R. Morales, A stretch of 17 amino acids in the prosaposin C terminus is critical for its binding to sortilin and targeting to lysosomes. J Histochem Cytochem 58, 287–300 (2010).

18. A. Darmoise, P. Maschmeyer, F. Winau, The immunological functions of saposins. Adv Immunol 105, 25–62 (2010).

19. T. Kolter, K. Sandhoff, Principles of lysosomal membrane digestion: stimulation of sphingolipid degradation by sphingolipid activator proteins and anionic lysosomal lipids. Annu Rev Cell Dev Biol 21, 81–103 (2005).

20. F. Ciaffoni et al., Saposin D solubilizes anionic phospholipid-containing membranes. J Biol Chem 276, 31583–31589 (2001).

21. F. Winau et al., Apoptotic vesicles crossprime CD8 T cells and protect against tuberculosis. Immunity 24, 105–117 (2006).

22. W. Jiang et al., The receptor DEC-205 expressed by dendritic cells and thymic epithelial cells is involved in antigen processing. Nature 375, 151–155 (1995).

23. J. Volckmar et al., Chemical Conjugation of a Purified DEC-205-Directed Antibody with Full-Length Protein for Targeting Mouse Dendritic Cells In Vitro and In Vivo. J Vis Exp, (2021).

24. I. J. Fidler, Biological behavior of malignant melanoma cells correlated to their survival in vivo. Cancer Res 35, 218–224 (1975).

25. S. A. Quezada et al., Limited tumor infiltration by activated T effector cells restricts the therapeutic activity of regulatory T cell depletion against established melanoma. J Exp Med 205, 2125–2138 (2008).

26. M. K. Ruhland et al., Visualizing Synaptic Transfer of Tumor Antigens among Dendritic Cells. Cancer Cell 37, 786–799 e785 (2020).

27. Y. Jiang et al., Prosaposin is a biomarker of mesenchymal glioblastoma and regulates mesenchymal transition through the TGF-beta1/Smad signaling pathway. J Pathol 249, 26–38 (2019).

28. S. Ishihara, D. R. Inman, W. J. Li, S. M. Ponik, P. J. Keely, Mechano-Signal Transduction in Mesenchymal Stem Cells Induces Prosaposin Secretion to Drive the Proliferation of Breast Cancer Cells. Cancer Res 77, 6179–6189 (2017).

29. S. Y. Kang et al., Prosaposin inhibits tumor metastasis via paracrine and endocrine stimulation of stromal p53 and Tsp-1. Proc Natl Acad Sci U S A 106, 12115–12120 (2009).

30. S. Wang et al., Development of a prosaposin-derived therapeutic cyclic peptide that targets ovarian cancer via the tumor microenvironment. Sci Transl Med 8, 329ra334 (2016).

31. M. Liu et al., TGF-beta suppresses type 2 immunity to cancer. Nature 587, 115–120 (2020).

32. L. Yuan, C. R. Morales, Prosaposin sorting is mediated by oligomerization. Exp Cell Res 317, 2456–2467 (2011).

33. X. Zhou et al., Impaired prosaposin lysosomal trafficking in frontotemporal lobar degeneration due to progranulin mutations. Nat Commun 8, 15277 (2017).

34. H. Salmon et al., Expansion and Activation of CD103(+) Dendritic Cell Progenitors at the Tumor Site Enhances Tumor Responses to Therapeutic PD-L1 and BRAF Inhibition. Immunity 44, 924–938 (2016).

35. S. Spranger, D. Dai, B. Horton, T. F. Gajewski, Tumor-Residing Batf3 Dendritic Cells Are Required for Effector T Cell Trafficking and Adoptive T Cell Therapy. Cancer Cell 31, 711–723 e714 (2017).

36. J. E. Jang et al., Crosstalk between Regulatory T Cells and Tumor-Associated Dendritic Cells Negates Anti-tumor Immunity in Pancreatic Cancer. Cell Rep 20, 558–571 (2017).

37. K. C. Barry et al., A natural killer-dendritic cell axis defines checkpoint therapy-responsive tumor microenvironments. Nat Med 24, 1178–1191 (2018).

38. P. Sharma et al., The Next Decade of Immune Checkpoint Therapy. Cancer Discov 11, 838–857 (2021).

## References and Notes

39. N. Fujita et al., Targeted disruption of the mouse sphingolipid activator protein gene: a complex phenotype, including severe leukodystrophy and wide-spread storage of multiple sphingolipids. Hum Mol Genet 5, 711–725 (1996).

40. I. S. Zeelenberg et al., Targeting tumor antigens to secreted membrane vesicles in vivo induces efficient antitumor immune responses. Cancer Res 68, 1228–1235 (2008).

41. W. Morelle, J. C. Michalski, Analysis of protein glycosylation by mass spectrometry. Nat Protoc 2, 1585–1602 (2007).

